# pomp-astic Inference for Epidemic Models: Simple vs. Complex

**DOI:** 10.1101/125880

**Authors:** Theresa Stocks, Tom Britton, Michael Höhle

## Abstract

Infectious disease surveillance data often provides only partial information about the progression of the disease in the individual while disease transmission is often modelled using complex mathematical models for large populations, where variability only enters through a stochastic observation process. In this work it is shown that a rather simplistic, but truly stochastic transmission model, is competitive with respect to model fit when compared with more detailed deterministic transmission models and even preferable because the role of each parameter and its identifiability is clearly understood in the simpler model. The inference framework for the stochastic model is provided by iterated filtering methods which are readily implemented in the `R package pomp`. We illustrate our findings on German rotavirus surveillance data from 2001 to 2008 and calculate a model based estimate for the basic reproduction number *R*_0_ using these data.

## 1 Introduction

Infectious disease epidemiology is concerned with the control and, potentially, the eradication of the disease from the population (Keeling and Rohani, 2008). During the last decades the growing field of mathematical modelling has provided essential tools for gaining understanding of infectious disease dynamics, monitoring public health data and early outbreak detection (Anderson and May, 1991; Keeling and Rohani, 2008; Diekmann et al., 2013). Moreover, mathematical models play a crucial role in infectious disease prevention by assessing the impact of different control measures, e.g. vaccination strategies, thus, developing the quantitative foundation for advising control policies, e.g. in global health (Heesterbeek et al., 2015).

Depending on which questions the modelling efforts are supposed to answer the resulting models can from a mathematical viewpoint be simple or complex. With an increasing number of model components it becomes more challenging to identify the role of each component and its interplay with the system (Keeling and Rohani, 2008). On the other hand, when developing models together with infectious disease specialists, a certain detail is required in order for a model to be considered as being realistic. It is exactly this trade-off between simplicity and “reality” which is the focus of this work. Inspired by May (2004) we investigate the question if a simple susceptible-infected-recovered (SIR) type model, in which the dynamics are clearly understood, is capable of explaining large population epidemiological data and, even more, if it is competitive with respect to model fit to a more detailed model. We illustrate our findings on rotavirus transmission in Germany, however, the modelling background could be easily modified to explain other infectious diseases which have comparable transmission characteristics, e.g. influenza, chickenpox or pneumococcus disease.

A crucial part of assessing a models usefulness is the calibration of model parameters from data, if adequate data are available. However, the number of model parameters that can be identified from data depends heavily on the quality and information contained in the available data and imposes a limit to the explanatory power of the model (Bretó et al., 2009). This is especially compelling for routine infectious disease surveillance data, because observations are recorded at discrete times and commonly describe just one aspect of the disease progression, e.g. the number of new infections aggregated over a certain time interval, say, one week. Event time data, such as infection and recovery times, are rarely available at a larger scale (O’Neill, 2010). Moreover, the disease dynamics are subject to various structural fluctuations such as seasonal and environmental changes, different genotypes or variability in social behaviour such as super spreaders etc. Most of these possible sources of variability are in part not sufficiently understood or not measurable in any way. Hence, stochastic variation is potentially an essential element to capture the unknown or non-measurable influences.

Models based on a deterministic transmission model component explain all variability in the data by an observational component since this is the only element where stochasticity enters, see e.g. in Weidemann et al. (2013); Althaus (2014). The predictive strength of these models is ambiguous since, in observed data, we see fluctuations which are clearly not due to variation in data collection alone, but due to phenomena not captured by the model. The sources of variability might be numerous and not even directly identifiable, but they all have in common that the underlying epidemic dynamics are subject to fluctuations which are independent of population size. The question arises if models that have much simpler features abstracting epidemiological insights, but address this structural variability, are competitive or even better suited to explain epidemiological data than their often very complex deterministic counterparts. These deterministic transmission models are very detailed with respect to disease states, age classes or spatial components, however, fail to address variability in the transmission model itself. The advantages of simpler models are clear: the dependence structure of parameters is easier to disentangle, the computational effort decreases with less detailed models and the information contained in data might not suffice to inform very complex transmission models. We investigate this question for rotavirus transmission in Germany between 2001 and 2008. This data set is particularly well suited for our investigations, since it was previously analysed in Weidemann et al. (2013) and Weidemann et al. (2014) as part of advising the German standing board of vaccination (STIKO) about the possible impact of a recommendation of rotavirus vaccination for children. This data set is hence a good use-case for comparing a very detailed deterministic transmission model, which explains variability of the data purely by observational noise, to a much simpler but stochastic transmission model including structural variability. We will follow the approach of Bretó et al. (2009) and include structural noise in form of stochastic transmission rates into our model. This has been used in previous studies of infectious diseases of malaria (Bhadra et al., 2011), measles (He et al., 2009), polio (Martinez-Bakker et al., 2015) and rotavirus (Martinez et al., 2016).

Our analysis will show that a simple model is capable to explain the above rotavirus data. However, it also clearly shows that it is important to include variability in the form of over-dispersion but also structural noise into the model.

### 1.1 Disease characteristics of rotavirus

Rotavirus is a childhood disease and the primary cause for gastroenteritis in infants and young children while adults are rarely infected (Dennehy, 2015; Grimmwood and Lambert, 2009). By the age of five nearly every child has been infected with the virus at least once (Bernstein, 2009). The virus spreads by direct transmission on the faecal oral route. The incubation time of rotavirus is around 2 days while severe symptoms last for approximately 4-8 days (CDC, 2016). Infected children may have severe watery diarrhoea, often accompanied by vomiting, fever, and abdominal pain (CDC, 2016). Children may develop rotavirus disease more than once, because neither natural infection with rotavirus nor rotavirus vaccination provides full immunity (protection) from future infections. Usually an individual’s first infection with rotavirus causes the most severe symptoms (CDC, 2016). Due to (partial) immunity acquired in childhood, most adults are hardly susceptible to rotavirus and if infected only show mild or no symptoms i.e. asymptomatic infections. However, a higher incidence is observed in elderly people above 60 years again, compare e.g. Figure 1, which might be due to a weaker immune system in higher ages which leads to an increased susceptibility.

**Figure 1:**
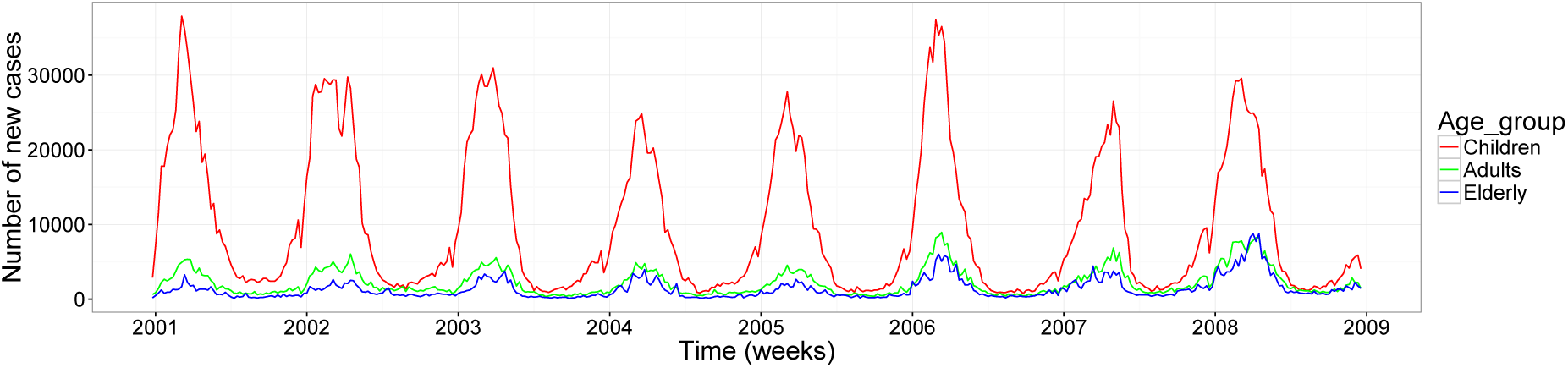
Weekly number of reported rotavirus cases among the three aggregated age groups of 0-4, 5-59 and 60-99 years of age in Germany 2001-2008 scaled by under-reporting factors inferred in Weidemann et al. (2013).

The data we analyze is the weekly reported number of new, laboratory-confirmed rotavirus cases among children, adults and elderly from 2001 until 2008 in Germany. Thereafter a significant impact on the rotavirus incidence by the increasing vaccination coverage is observed (Weidemann et al., 2013). The data are available through SurvStat@RKI (RKI, 2016). One problem with the available routine surveillance data is underreporting, however, since our focus is on methodological insights, we use the results from Weidemann et al. (2013) and simply scale up the available data with these inferred factors. The values we used for scaling up are the median estimates of the underreporting parameter from the averaged posterior distribution inferred in Weidemann et al. (2013), namely 4.3% in the former western federal states (WFS) and 19.0% in the former eastern federal states (EFS) between 2001-2004 and 6.3% in WFS and 24.1% in EFS from 2005 onward.

The time series depicted in Figure 1 are the original data scaled up by these inferred factors for three age groups, i.e. infants and younger children (up to 4 years of age), individuals between 5 and 60 years of age and older adults (60 years and older). The case report data clearly shows that the occurrence of rotavirus varies seasonally, peaking in March except in year 2007 where the season starts slightly later.

Mathematical modelling with SIR-type models of rotavirus has already been carried out for the USA (Pitzer et al., 2009), England and Wales (Atkins et al., 2012; Atchison et al., 2010; Pitzer et al., 2012), Western Europe (Van Effelterre et al., 2010), Australia (Shim et al., 2006), Kyrgyzstan (Freiesleben de Blasio et al., 2010), Germany (Weidemann et al., 2013) and Bangladesh (Martinez et al., 2016).

In the following, we address the question of how many parameters can be identified using the available data, demonstrate our modelling and inference framework and finally make some qualitative statements about the endemic rotavirus situation in Germany during that period by estimating susceptibilities of different age groups to the disease and the basic reproduction number. Conveniently, inference methods for both a deterministic and stochastic underlying transmission processes are implemented in the `R package pomp` (King et al., 2016). Since the package was developed fairly recently and spans a wide collection of inference tools we also comment on our experiences with the package and discuss its scientific as well as practical use. The present work is organized as follows. In Section 2 we present three different but related transmission models, namely a general stochastic SIR-type model, a stochastic SIR model with stochastic transmission rates and a limiting deterministic SIR model. We explain how the transmission models are connected to the data via an observational component and shortly present the inference framework for each model. Before carrying out the inference we mathematically investigate how many parameters are actually identifiable with the information the data contains and fix the other parameters at literature informed values. In Section 3 we present our inference results and compare the findings to the approach used in Weidemann et al. (2013). In Section 4 we discuss the gained knowledge, limitations of the model and conclusions.

## 2 Methods

In the following, we formulate a transmission model for rotavirus which is inspired by the work of Weidemann et al. (2013). It is simplified concerning the number of states and age classes but includes more variability than just over-dispersion by allowing for structural noise in the transmission rates.

### 2.1 Partially observed dynamical system

One practical way to answer infectious epidemiology related questions in a model and data driven way is the use of partially observed dynamical systems (Ionides et al., 2006). These models explain how data and a disease transmission model are related as they consist of two components: an unobserved, time-continuous state process, which operates on the population level describing the dynamics of disease spread, and an observation model, which describes how the data collected at discrete points in time is connected to the transmission model. The transmission model can be either deterministic or stochastic, while observations are most often assumed to be stochastic, centered around an adequate summary of the transmission model. If the transmission model is assumed to be a Markov process then the system is called a partially observed Markov process (short: pomp) (King et al., 2016). Figure 2 illustrates the components of a partially observed dynamical system.

**Figure 2:**
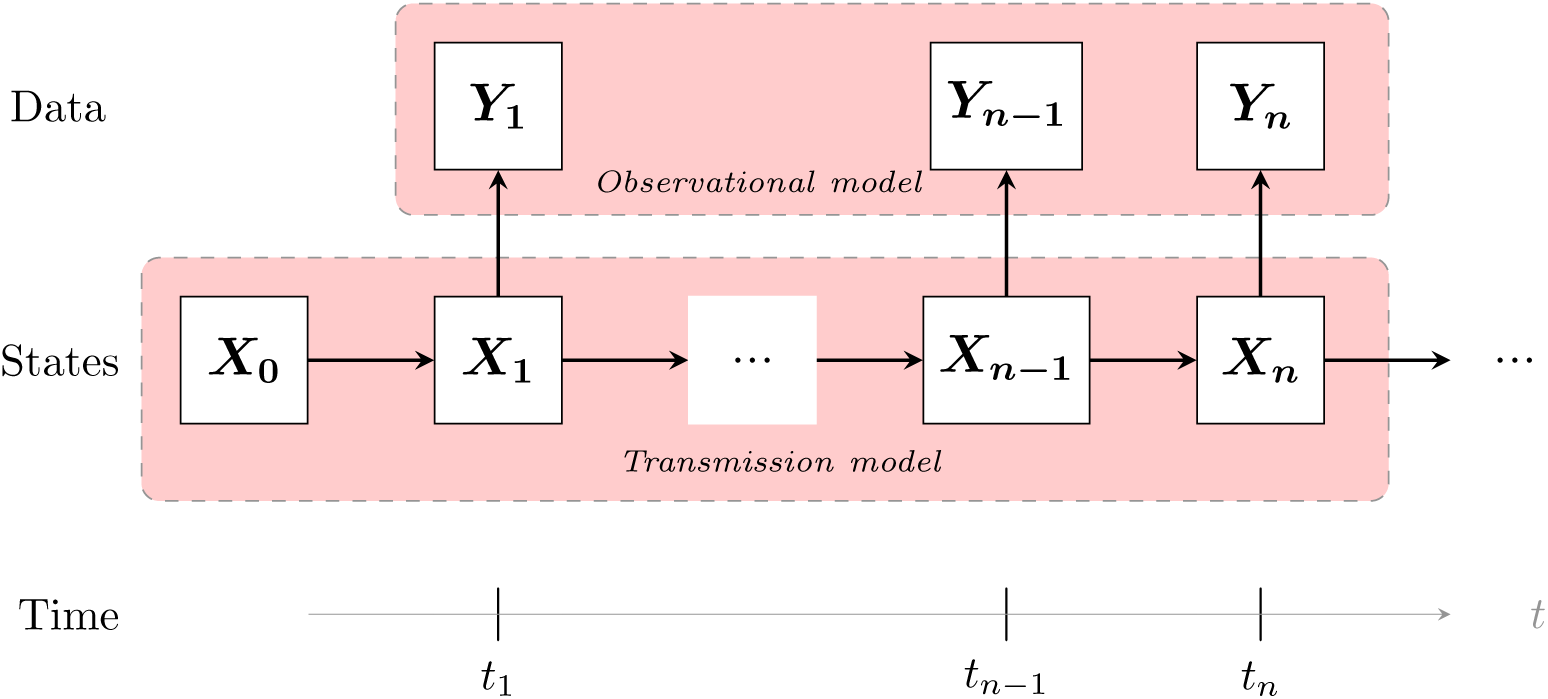
A partially observed dynamical system where ***Y**_i_, i* = 1, …, *n* denotes the observations at time *t_i_*, which depend on the state of the transmission process ***X**_i_* at that time.

### 2.2 Stochastic transmission model

We assume that the transmission model is a Markov process where individuals move between compartments at random times, as described in detail in what follows.

The schematic representation of our model is given in the flow diagram of Figure 3 where arrows indicate the movement between the compartments. We subdivide the population into nine discrete compartments, stratified by the age and health status of the individuals. Although age is time continuous,a compartmental approach is often adequate because children are generally grouped into daycare or school classes of a given age cohort (Keeling and Rohani, 2008). We consider a model that subdivides the population into three age classes, identified by the indices 1, 2 and 3 respectively. We choose this age stratification because the disease burden is highest for young children, very low for children over 5 years of age and rises again later in life. The variables *S_k_*(*t*), *I_k_*(*t*), *R_k_*(*t*) ∈ ℕ with *k* ∈ {1, 2, 3} count the number of susceptible, infectious, and recovered in each of the three age groups at time *t* ∈ ℝ^+^.

**Figure 3:**
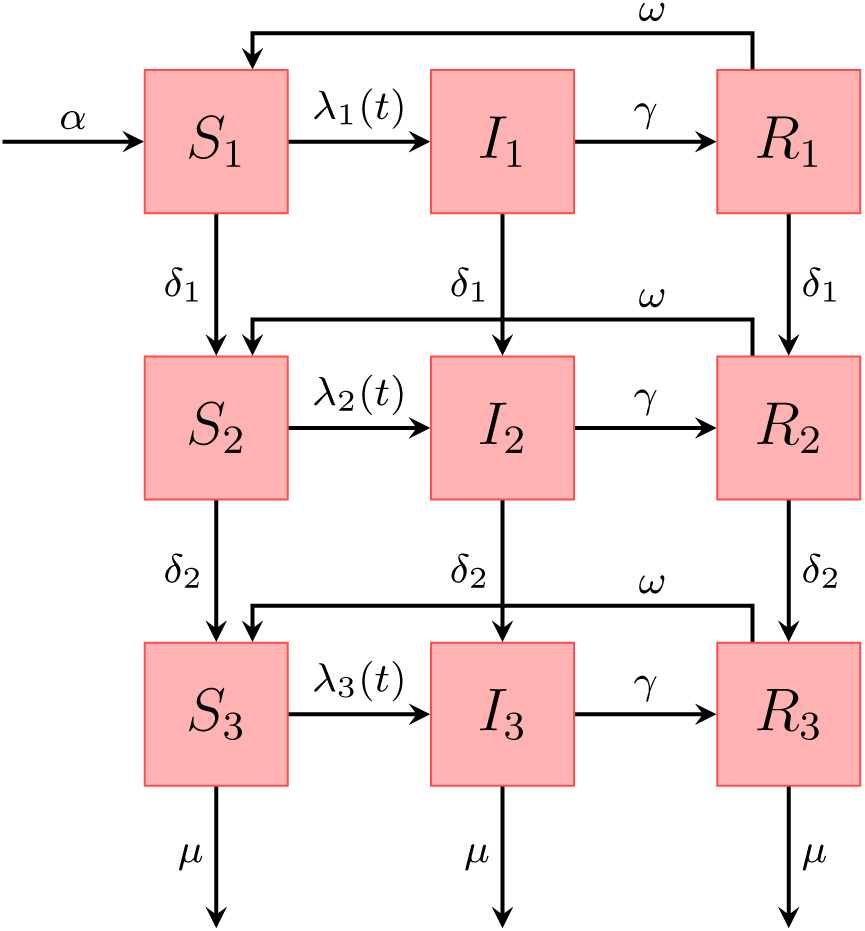
Schematic representation of the states in our SIRS model with three age classes. The rates represented by the arrows are explained in the text

Concerning the rates, children “age” with a rate *δ*_1_ into the second (=adult) age class and adults “age” with a rate *δ_2_* into the class of elderly. The vector **λ**(*t*) = (λ_1_(*t*), λ_2_(*t*), λ*_3_*(*t*))′ is called the *force of infection* and is the per capita rate at which susceptible individuals get infected which depends on the number of currently infected individuals *I_k_*(*t*). Susceptible individuals can become infected by transmission from an infectious individual in one of the three age groups. The recovery rate *γ* ∈ ℝ^+^ is assumed to be independent of age. For reasons of parameter identifiability, which will be discussed in detail later, we also assume that waning of immunity is independent of age and immunity from infection lasts for a limited, exponentially distributed period with mean 1/*ω* ∈ ℝ^+^ after which the individual is again fully susceptible.

Since the observed disease dynamics evolve over several years, the model has to account for demographics. We assume that the average population size *N* is constant, which implies that the birth rate *αN* ∈We assume that the average population ℝ^+^ equals the overall death rate. Furthermore, it is assumed that death can only occur in the last age class with rate *μ* ∈ ℝ^+^ independent of the health status of the individual. This seems reasonable because in developed countries 90 % of the mortality comes from individuals older than 60 years, hence premature death can be ignored for our purposes (Atkins et al., 2012). To keep the population size constant on average it is assumed that *α* is population independent and equals the inverse of the average lifetime and *μ* equals the inverse of the average time spend in the last age-class. In the following, the notation is adopted from King et al. (2016). We let *N_AB_*(*t*) denote a stochastic counting process which counts the number of individuals which have moved from compartment A to compartment B during the time interval [0, *t*) with *A, B*∈ *χ*, where *χ* = {*S*_1_, *S*_2_, *S*_3_, *I*_1_ *I*_2_, *I*_3_, *R*_1_, *R*_2_, *R*_3_} contains all compartments of our model. Furthermore, *N*_▪_*_A_* (*t*) counts the number of births and *N*_▪_*_A_* (*t*) counts the number of deaths in the respective compartment up until time *t*. The infinitesimal increment probabilities of a jump between compartments connected by an arrow (Figure 3) fully specify the continuous time Markov process describing disease transmission. Let ∆*N_AB_*(*t*) = N*_AB_*(*t* + *τ*) − *N_AB_*(*t*) count the number of individuals changing compartment in an infinitesimal time interval *τ* > 0. Then we define for the model depicted in Figure 3 the following system of transition rates:

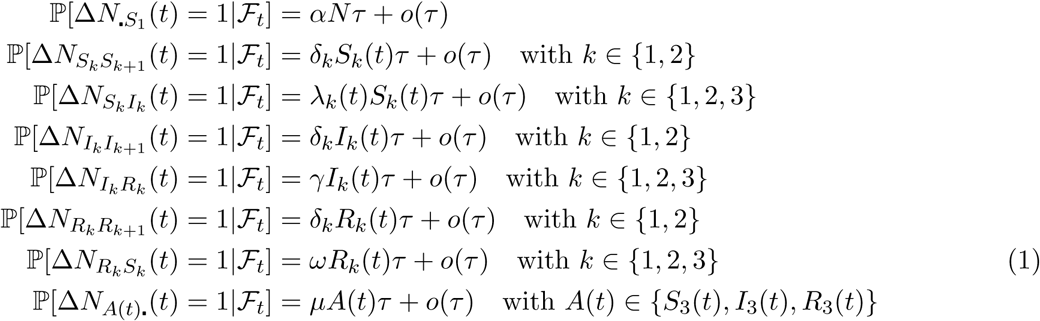
with the filtration 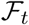 = {*S*_1_(*u*), *I*_1_(*t*), *R*_1_(*u*), *S*_2_(*u*), *I*_2_(*u*), *R*_2_(*u*), *S*_3_(*u*), *I*_3_(*u*),*R*_3_(*u*), ∀ 0 ≤ *u* ≤ t} denoting the history of the process until time *t*. The transmission rates are related with the state variables in the following way:

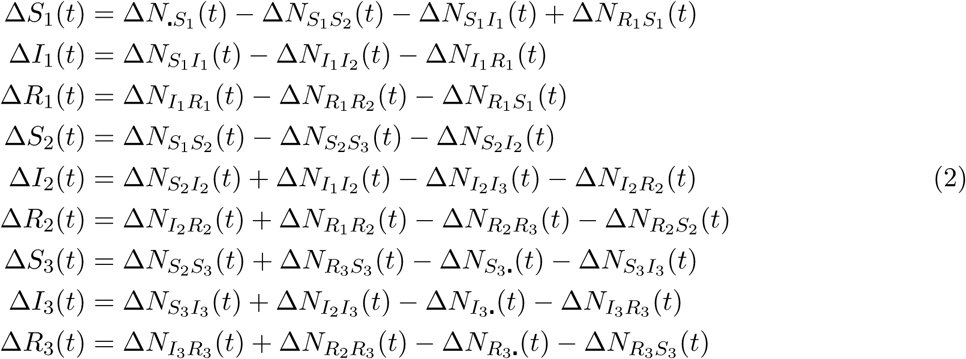

A consequence of the above model formulation is that we assume that individuals change from one compartment to another according to an exponential distribution. The number of newly infected individuals in age class *k* accumulated in each observation time period [*t_j_, t_j_*_+1_) is then given as

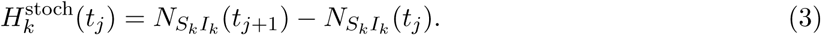

The force of infection, written in vector notation, consists of the following components

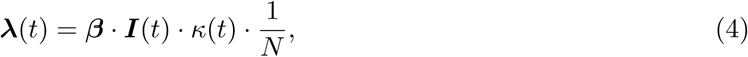
where ***I***(*t*) = (*I*_1_(*t*), *I*_2_(t), *I*_3_(*t*))′. Disease transmission is represented in a transmission-matrix ***β*** = (*β _kj_*)*_k,j_*_∈{1,2,3}_, which is often called a WAIFW (who acquires infection from whom)-matrix (Keeling and Rohani, 2008). The parameter *β_kj_* denotes the average number of infectious contacts of infected individuals of age group *j* with susceptible individuals of age group *k* per time unit and hence is the product of contact rates and transmission probability. Our modelling should account for the possibility that individuals in each age class *k* could have a different immune status due to age and hence having differing susceptibility to the disease. For this work we assume age dependent infectious contact numbers such that *β_kj_* = *β_k_* for all *k*,*j* ∈ {1, 2, 3}. Furthermore, motivated by the rotavirus application, we introduce a seasonal periodic forcing because we assume that those pronounced fluctuations are likely due to social aggregations of the host like in daycare institutions and schools which are closed during summer or climate changes. In contrast to ***λ***(*t*)which is state dependent, the seasonal forcing function *κ* (*t*) is assumed to be purely time dependent i.e. it has the same effect in all age groups. We let

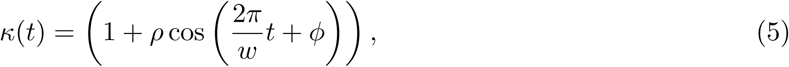
where *ρ* ∈ [0, 1] is the amplitude of the forcing, 2*π*/ω ∈ ℝ^+^ is the period of the forcing (e.g. if the time unit is weeks, then *ω* = 52) and *ϕ* ∈ [0, 2*π*] is the phase shift parameter. Note that with the choice of forcing in equation (5) the parameter *β_k_* denotes the baseline or average transmission rate of an individual of age group j with age group k which varies between (1 − *ρ*)*β_k_* and (1 + *ρ*)*β_k_* during the year (Keeling and Rohani, 2008).

#### Extension: structural noise

Including sufficient stochasticity in a model as a way to capture drivers and phenomena not covered otherwise by the model (e.g. late season start) is essential if one wants to assess the predictive power of the model (Bretó et al., 2009). So far, we have accounted for stochasticity in the underlying system by assuming that individuals move between classes at random times. However, for large population sizes the stochastic system approaches the deterministic system and, hence, the role of randomness diminishes as the population size increases. The same occurs by modelling disease spread via stochastic differential equations (Fuchs, 2013). One way to introduce variability, which is independent of population size, is to assume stochastic fluctuations in the transmission rates. In this work, we will follow the approach of Bretó et al. (2009) and introduce a time continuous stochastic process *ξ*(*t*) which fluctuates around the value one and is multiplied onto the transmission rate. It can be shown that by choosing the corresponding integrated noise process Γ(*t*) in a way such that its increments are independent, stationary, non-negative and unbiased the Markov property is retained (Bretó et al., 2009). One convenient example for a process which satisfies these conditions is a Lévy process with

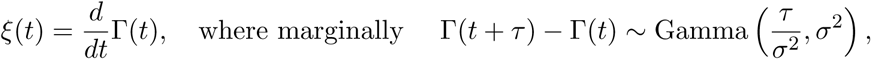
and where τ/*σ*^2^ denotes the shape and *σ*^2^ the age-independent scale parameter with corresponding mean *τ* and variance *τ*σ**^2^. Note, that the integral of *ξ*(*t*) over a time interval is well defined even though the sample paths of Γ(*t*) are not formally differentiable (Karlin and Taylor, 1981; Bretó et al., 2009). The parameter *σ*^2^ is called the *infinitesimal variance* parameter (Karlin and Taylor, 1981). We build this into the model by letting the force of infection be

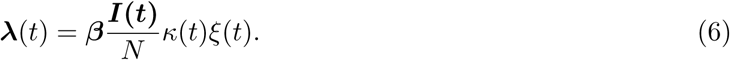

### 2.3 Deterministic transmission model

Calculating the expected values of the equations in (2)with the aid of (1), dividing by *τ*, and taking the limit as *τ* → 0, we obtain the underlying deterministic form of the three age strata model:

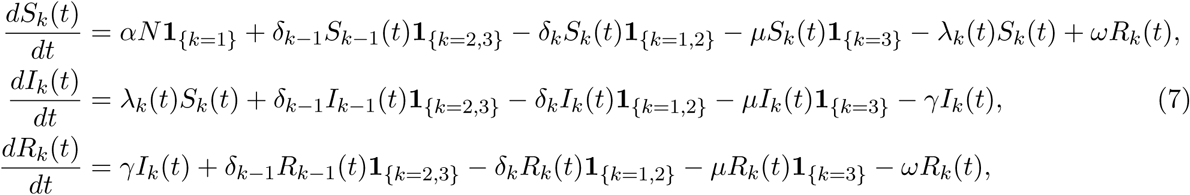
for *k* = 1, 2, 3 and initial values satisfying 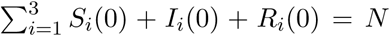. The number of newly infected individuals in age class k accumulated in each observation time unit [*t_j_, t*_*j*+1_) is then given as

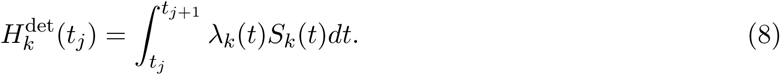

#### Analytical solution of the deterministic transmission model without seasonality

Optimally, all parameters of a model can be inferred from the data, however, as we only observe an aggregation of a part of the system at discrete times we would like to investigate which and how many parameters of our model are actually identifiable from data. We note that the seasonal component in the data allows us to estimate the phase shift parameter *ϕ* and the amplitude of the forcing *ρ*, however, there is no closed form solution for those two parameters. What remains to be investigated is how many parameters can be estimated from the data without this seasonal component which corresponds to analyzing the system in endemic state. For this purpose, we divide the ODE system in (7) by *N* so we obtain the fractions of the population being susceptible, infected and recovered, i.e. *s_k_*(*t*) = *S_k_*(*t*)/*N, i_k_*(*t*) = *I_k_*(*t*)/*N* and *r_k_*(*t*) = *R_k_*(*t*)/*N, k* ∈ {1, 2, 3} with the sum over all fractions adding up to one and write *λ_k_*(*t*) = *β_k_i*(*t*) with 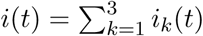. We are interested in the system at equilibrium, i.e. when *ds*_*k*_/*dt* = *di_k_*/*dt* = *dr_k_*/*dt* = 0. Here, we work out the values of the variables which we denote by 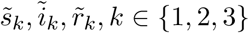 (Keeling and Rohani, 2008).

In order to obtain an artificial endemic state without seasonality we take for each age group the mean over time and treat the obtained values as our data *Y_k_*(*t_j_*) which is the number of newly reported rotavirus cases aggregated over reporting intervals [*t_j_, t_j_*+_1_), *j* ∈ {0, 1, 2, …} in age class *k* ∈ {1, 2, 3}. In the endemic state 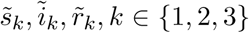 also the number of reported cases is in equilibrium, hence *Y_k_*(*t*_j_) = *Y_k_*(*t*_j_+_1_) for all *j* ∈ {0, 1, 2, …}. For *t_j_*_+1_ − *t_j_* = 1 and assuming no observational noise, this translates mathematically to

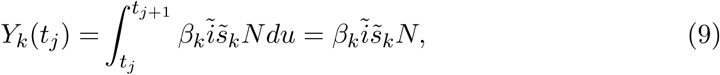
with 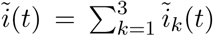, and which is independent of the time point *t_j_*. Hence, we will only write *Y_k_* := *Y_k_*(*t*_j_) in what follows. Plugging (9) into the transformed equations (7) at equilibrium we obtain

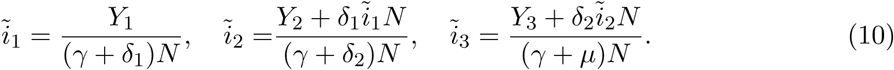

We conclude that the data thus can inform the three endemic states 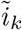, *k* ∈ {1, 2, 3}, if we assume that the demographic parameters for *μ, δ*_1_, *δ*_2_ as well as the recovery rate *γ*, are fixed and known. Furthermore, since in the deterministic model the total population size *N* as well as the population sizes *N_k_* = *S_k_*(*t*) +*I_k_*(*t*) + *R_k_*(*t*) of age class *k* are constant, we know that

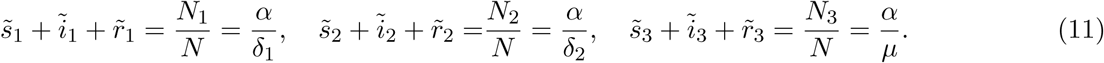

These equalities follow from the fraction of the population in age class *k* being equal to the fraction of the average lifetime spend in each age class. We can therefore express 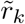 in terms of 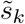 and 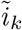, if we assume additionally that the birth rate parameter *α* is fixed and known. The variables that remain to be estimated are 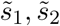 and 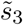. It can now be seen that by further assuming *ω* to be fixed and known we can also estimate *β*_1_, *β*_2_ and *β*_3_. For this, we equate the above system to zero and solve the equations. We obtain by analytical derivations

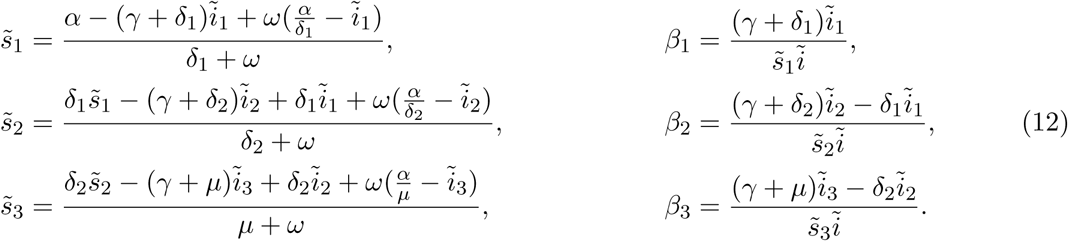

We thus conclude the following: by assuming the demographic parameters α, *μ, δ*_1_, *δ*_2_, the recovery rate *γ* and the immunity waning rate *ω* to be fixed and known it is possible to estimate for *φ, ρ, β*_1_, *β*_2_ and *β*_3_ from the available data. In case that more parameters should be estimated from the data additional sources of information are needed.

### 2.4 Calculation of *R*_0_

An important mathematical characteristic of an epidemic model is its basic reproduction number *R*_0_. It is the expected number of new infections by a typical infectee during the early stage of an epidemic when everyone is susceptible (Andersson and Britton, 2000). We can calculate the basic reproduction number *R*_0_ for the deterministic transmission model which is, also, representative for the stochastic transmission model, because it approaches the deterministic system for a large population size *N*. Hence, let *π_k_* = *N_k_*/*N* denote the community fraction of age class *k* ∈ {1, 2, 3} in the population and *ν_k_* = 1/(*γ* + *δ_k_*) for *k* = 1, 2 or *v_k_* = 1/(*γ* + *μ*) for *k* = 3 the average length an individual of type *k* ∈ {1, 2, 3} stays in the infectious compartment-note that the infectious period is reduced due to some individuals ageing or dying while infectious Keeling and Rohani (2008). With the seasonal forcing function chosen as in equation (5) the expected number of *k* individuals a *j* individual infects if everyone is susceptible is *β_k_ π_k_ν_j_*, which represents a yearly average. The matrix

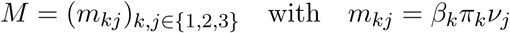
defines the expected number of new infections of individuals of a certain age caused by individuals of a certain age if everyone is susceptible. The basic reproduction number *R*_0_ can then be calculated as the largest eigenvalue of this matrix Andersson and Britton (2000), pp. 51-62. However, if *m_k_j* = *p_k_n_k_ν_j_* (i.e. “proportionate mixing” Diekmann et al. (2013), p. 176) the basic reproduction number can be written as

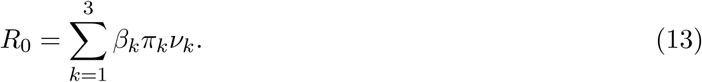

This yearly average will vary between (1 − *ρ*)R_0_ and (1 + *ρ*)*R*_0_ during the year.

### 2.5 Observation model

To incorporate the count nature of the observations a natural first assumption would be to model the reported cases as realizations of a Poisson distributed random variable with a given time dependent mean. However, the data suggests that the sample variance is larger than the sample mean, i.e. there is indication of over-dispersion in the data. A better choice in this case is therefore the negative binomial distribution which allows for additional variance.

Let the number of recorded cases *Y_k_*(*t_j_*), *k* ∈ {1, 2, 3} within a given reporting interval [*t_j_, t_j_*+_1_), *j* ∈ {0, 1, 2, …;} be

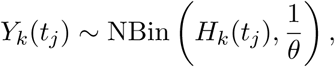
with *H_k_* (*t_j_*) being the true number of accumulated incidences in age class *k* per time unit [*t_j_, t_j_*+_1_) in the model, cf. equation (3) for the stochastic model and (8) for the deterministic model. Here NBin(*μ*, 1/*θ*) denotes the negative binomial distribution with mean *μ* and variance *μ* + *θμ*^2^. To reduce the number of parameters, the same dispersion parameter *θ* for all age classes is chosen as in Weidemann et al. (2013).

### 2.6 Inference and Implementation

#### 2.6.1 Simulation from the stochastic model

To generate realizations from the stochastic transmission model in (2) we can use the Gillespie algorithm (Gillespie, 1977). Given the current state of the system, the algorithm simulates the waiting time for every possible next event given its current state. Then it chooses the shortest waiting time and updates the number of individuals in each compartment and the overall time is incremented accordingly. The whole procedure is then repeated until a pre-defined stopping time is reached. The simulation of every individual event gives us a complete and detailed history of the process, however, it is usually a very time-consuming task for systems with large population and state space, because of the enormous number of events that can take place (Gillespie, 2001).

As a way to speed up such simulations we choose an approximate simulation method, the so called *τ-leap algorithm* which is based on the Gillespie algorithm. It holds all rates constant in a small time interval *τ* and simulates the numbers of events that will occur in this interval, then updates all state variables, computes the transition rates again and the procedure is repeated until the stopping time is reached (Erhard et al., 2010; King et al., 2015). Given the total number of jumps, the number of individuals leaving any of the states by the routes indicated by arrows (cf. Figure 3) during a time interval *τ* is then multinomially distributed (King et al., 2015). This is because Poisson random variables *X*_1_, …, *X_n_* conditioned on their sum, i.e. 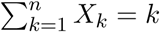, result in a multinomial distributed random variable with size parameter *k* and the transition probabilities proportional to their respective rates. Under appropriate conditions this procedure can produce significant gains in simulation speed with acceptable loss in accuracy (Gillespie, 2001).

#### 2.6.2 Likelihood for partially observed dynamical systems

Let ***Y****_n_* = (*Y*_1*n*_, …, *Y*_3*n*_)′ denote the random variable counting the observations at time *t_n_* in each of the three age-classes depend on the state of the transmission process ***X****_n_* = (***S***(*t_n_*), ***I***(*t_n_*), ***R***(*t_n_*)) at that time where, e.g. ***S***(*t_n_*) = (*S*_1_(*t_n_*), *S*_2_(*t_n_*), *S*_3_(*t*_3_))′, cf. Figure 2. Furthermore, we denote 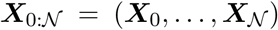 and the parameter vector by ***ψ*** The joint density of the states and the observations is then defined as the product of the one-step transmission density, *f*/_***X**n*_|_***X**n*−l_(***x**_n_*|***x***_*n*−1_; ***ψ***), the observation density, *f****Y***_n_|***X***_n_(***y***_*n*_|***x**_n_*; ***ψ***), and the initial density *f*_***X***0_(***x***_0_; ***ψ***) as

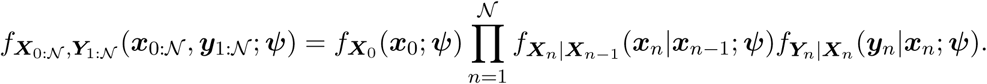

The likelihood of the parameter vector can then be written as the marginal density for a sequence of observations, 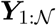 evaluated at the data, 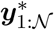 as

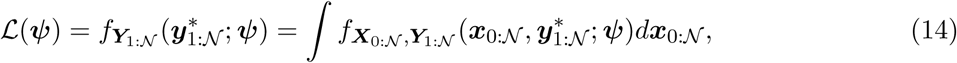
compare e.g. King and Ionides (2016a). Note that for our model this is a high-dimensional integral of dimension 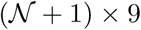, which is hard if not impossible to solve by any analytical means, see Section 2.6.4.

#### 2.6.3 Maximum likelihood estimation for determistic transmission model

Inference for partially observed dynamical systems with a deterministic underlying transmission model is relatively straightforward, because eq. (14) is computable. This is because the likelihood of observations depends only on the solution of the deterministic system (cf. (7) and (8)). More precisely, ***X**_n_* = ***x**_n_*(***ψ***) is a known function of ***ψ*** for each *n*. What is generally not known is the initial density *f***X**_0_(***x***_0_; ***ψ***), however we overcome this by initializing the system well ahead of the first observations such that simulations will equilibrate before the first observation is made. Hence, we choose the estimates from equations (10), (11) and (12) as the initial distribution and start the system 6 years prior to our first observation. The solution of the ordinary differential equation system can then be calculated numerically with, e.g., Runge-Kutta methods (Press et al., 2007). Given this solution, maximum likelihood estimation boils down to a classical numerical optimization problem for a non-linear function of the parameter vector.

##### Implementational details

Maximum likelihood estimation for partially observed dynamical systems with a deterministic underlying transmission model is, e.g., implemented in the `traj.match` function in the `R package pomp` (King et al., 2016) which accesses `R’s optim` function. As optimizing algorithm we chose the Nelder-Mead method Nelder and Mead (1965). In order to address the potential problem of local maxima in the optimization we use 100 randomly chosen parameter constellations as starting values for the fitting algorithm. These values are drawn uniformly from a hypercube which contains all sensitive parameter values. In general, if the inference procedure gives consistent results for starting values drawn at random from a hypercube this indicates that a global maximum has been found and a reliable global search has been performed King and Ionides (2016b). One practical problem for the models at hand is that if the relative convergence tolerance in the Nelder-Mead algorithm is very small, estimation fails due to a degeneracy of the Nelder-Mead complex. To get around this, we choose the relative tolerance as small as possible so the algorithm does not fail and re-use the obtained estimates as starting values for a second run of the inference algorithm with the same tolerance. Surprisingly, the implementation of the algorithm then usually takes a few additional iterations. This way we make sure that the algorithm really converges. Based on the obtained maxima for all starting values we select the one with the highest likelihood as our maximum likelihood estimator (MLE). To determine a 95% confidence interval (CI) for the obtained estimates we calculate the profile log-likelihood for each parameter of interest and invert Wilk’s (likelihood ratio) test to get the desired intervals Held and Sabanés Bové (2013). To construct a 95% pointwise prediction interval (PI) for model realizations we calculate the 2.5% as well as the 97.5% quantile of the respective response distribution at each observation time *t_j_* with mean 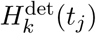 (eqn. (8)) using plug-in of the MLE and hence ignoring uncertainty in the parameters.

### 2.6.4 Maximum likelihood estimation for stochastic transmission model

For a partially observed stochastic transmission model the likelihood is not tractable, because knowledge of the parameters does not uniquely determine the solution of the transmission model and marginal likelihoods are very computationally demanding. During recent years different methods have been developed to overcome this problem. With increasing computational power, simulation-based methods gained more and more attention as well as modified likelihood-based approaches such as iterated filtering Ionides et al. (2011), simulated moments Kendall et al. (1999), synthetic likelihood Wood (2010), non-linear forecasting Sugihara and May (1990). Another track have been Bayesian approaches such as approximate Bayesian computations Toni et al. (2009); Liu and West (2001) and particle MCMC Andrieu et al. (2010). A good survey of such and other inference methods can be found in O’Neill (2010). In this work we will use iterated filtering which is a simulation and likelihood-based method which uses trajectories generated by the underlying transmission model as the basis for inference Bretó et al. (2009). To evaluate the likelihood of a partially observed Markov model for a set of parameters, the standard approach is to approximate the integral in equation (14) by Monte Carlo methods Robert and Casella (2004). However, it turns out that this method would not be very efficient because the way trajectories are proposed is completely unconditional of the data which leads to very imprecise estimates. A more efficient alternative is to factorize the likelihood in eq. (14) as

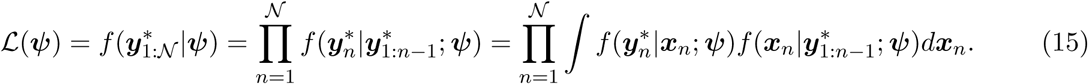

This formulation serves as basis for particle filter methods which efficiently use re-sampling techniques, details about which can be found in Doucet et al. (2001).

#### Implementational details

Iterated filtering as introduced by Ionides et al. (2006) and improved in Ionides et al. (2015) explores the parameter space by adding noise to the parameters of interest and at each iteration calculates the likelihood of the perturbed model by evaluating the particle filter. The algorithm is implemented in the `mif2` function in the `R package pomp` King et al. (2016). As the iterations proceed the intensity of the perturbation is successively reduced (“cooling”) and the loglikelihood of the perturbed model gradually approaches the loglikelihood of the model of interest. However, for a finite number of iteration steps, the loglikelihoods of the two models are not identical and a particle filter evaluation of the `mif2` model output using equation (15) is necessary. For the fitting carried out in the following we use 20 starting values drawn uniformly from a hypercube covering reasonable parameter values. To make calculations feasible with respect to time we accept approximation errors of the underlying stochastic transmission process by choosing a rather large time step size of 1/10 in the *τ* leaping algorithm, see Section 2.6.1. For the following inference procedure we use `Nmif=300` iterations, `Np=5000` particles, a cooling of the perturbations of `cooling.fraction.50=0.5` and random walk standard deviations `rw.sd` which vary dependent on the parameter between 0.001 and 0.2. For details see the pointed out manual.

For each of the 20 `mif2` outputs we run 10 particle filters, each with 1000 particles. From this we calculate the estimated mean of the loglikelihood (LL) and the standard error of the Monte Carlo approximation for every parameter set. Consequently, we choose the parameter constellation of the 20 possible with the highest loglikelihood as the maximum likelihood estimator. To obtain the 95 % confidence intervals for the parameters we construct the profile likelihood of each parameter and use Wilk’s test. This might seem overcomplicated at first sight since, intuitively, the parameter swarm should contain some measure of uncertainty. However, due to particle depletion the information about the local shape of the likelihood surface contained in the particles is not very reliable. It turns out that the profile likelihood is much more robust since it relies on multiple independent `mif2` computations. The 95% prediction intervals for the model realizations are calculated out of 1000 realizations of the model when using plug-in of the MLE, ignoring any uncertainty. To evaluate the model fit we investigate how often the data does not lie in this interval, for the transmission model as well as the whole model including the observational model. The implementation of the following calculations are available at Stocks (2017).

## 3 Results

In the following we will compare the model fit of four different models which are based on the same conceptual modelling of rotavirus transmission (cf. Figure 3) but differ with respect to the handling of stochasticity. We differentiate between deterministic and stochastic transmission models and consider stochastic observation processes with and without overdispersion. Model DtSt captures the least stochasticity and Model St+St+ the most, for model details compare Table 1. Our practical experience shows that for a stochastic model without overdispersion the iterated filtering algorithm fails, because the model is not variable enough to explain the data. Hence, we do not carry out inference for a stochastic transmission model with Poisson observations.

**Table 1:** Specification of the four investigated models.

| Model name | Transmission model | Observational model |
| --- | --- | --- |
| DtSt | deterministic | Poisson |
| DtSt+ | deterministic | negative binomial |
| StSt+ | stochastic | negative binomial |
| St+St+ | stochastic with structural noise | negative binomial |

As a first inference step we calculate the *β_k_*’s from the analytical solution given in Section 2.3. To demonstrate the performance of the methods used, we carry out a simulation study for each model first. In the second step we will use the insights gained to fit the four models to the observed rotavirus time series described in Section 1.1. In the previous section we found that only the susceptibility parameters *β_k_*, with *k* ∈ {1, 2, 3} and the parameters for the seasonal forcing function *ρ* and *φ* can be estimated from the available data. Moreover, we will estimate the overdispersion parameter *θ* for the Models DtSt+, StSt+ and St+St+ and additionally the shape parameter of the structural noise *σ*^2^ for Model St+St+. All other parameters are fixed at biological plausible values shown in Table 2. The inverse of the yearly birthrate equals the sum of the inverses of the yearly aging and death-rates which add up to 78.9 which is the total life expectancy at birth in Germany averaged between the years 2001 to 2008, taken from The World Bank (2016). The averaged population size *N* in Germany during that time period is taken from Statistisches Bundesamt (2016) and the reporting interval is one week. As in Atkins et al. (2012) we assume that all individuals are immune against rotavirus for an average of one year, after which they return to full susceptibility. A more detailed discussion on the choice of this assumption can be found in Section 4.

**Table 2:** Full list of notation and parameter values used in the model.

| Parameter | Explanation | Value | Unit |
| --- | --- | --- | --- |
| $N$ | population size | 82 372 825 | individuals |
| $\alpha$ | individual birth rate | $1/(52 \cdot 78.86912)$ | $\text{week}^{-1}$ |
| $\delta_1$ | ageing from age group 1 (age 0-4) to 2 (age 5-59) | $1/(52 \cdot 5)$ | $\text{week}^{-1}$ |
| $\delta_2$ | ageing from age group 2 (age 5-59) to 3 (age 60+) | $1/(52 \cdot 55)$ | $\text{week}^{-1}$ |
| $\mu$ | death rate (from age group 3 (age 60+) ) | $1/(52 \cdot 18.86912)$ | $\text{week}^{-1}$ |
| $\beta_k$ | susceptibility of age group $k \in \{1, 2, 3\}$ | to be estimated | individuals/week |
| $\omega$ | immunity waning rate | $1/(1 \cdot 52)$ | $\text{week}^{-1}$ |
| $\gamma$ | recovery rate | 1 | $\text{week}^{-1}$ |
| $\phi$ | phase shift of the seasonal forcing | to be estimated | 1 |
| $\rho$ | amplitude of the seasonal forcing | to be estimated | 1 |
| $\lambda_k(t)$ | force of infection of age group $k \in \{1, 2, 3\}$ | defined by eq. (4) | individuals/week |
| $\kappa(t)$ | seasonal forcing function | defined by eq. (5) | 1 |
| $\theta$ | overdispersion parameter | to be estimated | 1 |
| $\sigma^2$ | scale parameter of the Γ-noise | to be estimated | 1 |
| $\tau$ | time step | to be estimated | week |
| $R_0$ | basic reproduction number | to be estimated | individuals |

### 3.1 Analytical solution of *β_k_*

Our investigations showed that the inference algorithms are very sensitive to starting values of the parameters. Hence, we use the analytical solution of the deterministic transmission model without seasonality from Section 2.3 as a first step. We then use the obtained estimates for *β_k_*, as starting values for the other estimation procedures of all four models and the endemic state estimates as initial values for the transmission model.

In the rotavirus data the mean number of weekly reported new cases in each age group between the years 2001 and 2008 are 10009, 2254, and 1364 for children, adults and elderly respectively. Solving the calculations from equation (12) we obtain

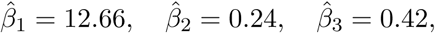
for the deterministic system. This supports what was stated initially that children are highly susceptible to the disease while adults enjoy a higher protection. However, with higher age the susceptibility increases slightly again. Moreover, calculating the basic reproduction number it follows from equation (13) that 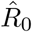. One could argue that this value is surprisingly close to the value one for a highly infectious disease as rotavirus. It should, however, be noted that the susceptibility for children is very high, but children in our model only make up approximately 6% of the total population. That the susceptibilities of the two older age groups are this low depends on the assumption of an average length of immunity of one year in all age classes. One possible explanation is that this is not adequate and that immunity lasts longer or under-reporting is higher in the older age classes which both leads to higher values for *R*_0_. We refer to Section 4 for a more detailed interpretation of these findings.

### 3.2 Simulation Study

In this section we perform a simulation study to demonstrate the correctness of the inference method for each of the four models. For this we generate one realization of each model with parameters chosen as the analytical solution, which we then treat as data to estimate parameters from. Note, that since the data is different for each simulation study the loglikelihoods of the four models are not directly comparable. After the inference we compare the obtained estimates to the true parameters which serves as validation of our implementation. The results can be found in Table 3. For all models the estimated parameters are in good accordance with the true parameters. As an example, Figure A.9 shows the simulated data from Model DtSt+, together with pointwise 95% prediction intervals obtained from the model when using plug-in of the MLE for the solution of the ODE. In order to check if observational noise and structural noise are distinguishable and how the estimation results change under model miss-specification we carried out a robustness study (not shown here). For this we generated one realization of each model and additionally a realization of a model with structural noise and Poisson observations and fitted the obtained realizations to every model respectively. We found that both noise components are indeed distinguishable from each other and parameters are estimated correctly even under model miss-specification.

**Table 3:** Simulation study results for the models with starting interval (SI), true parameter (TP), maximum likelihood estimator (MLE) and calibration of the 95% confidence interval (CI). Moreover, we report loglikelihood (LL) plus/minus the standard error of the Monte Carlo approximation of the loglikelihood.

| **Model DtSt** (LL: -6 517.864) |  |  |  |  |  |  |
| --- | --- | --- | --- | --- | --- | --- |
| | $\beta_1$ | $\beta_2$ | $\beta_3$ | $\rho$ | $\phi$ | |
| SI | [12.000, 16.000] | [0.200, 0.400] | [0.300, 0.500] | [0.120, 0.160] | [0.00, 0.200] |  |
| TP | 12.657 | 0.238 | 0.420 | 0.150 | 0.100 |  |
| MLE | 12.657 | 0.238 | 0.420 | 0.150 | 0.100 |  |
| CI | [12.651, 12.664] | [0.238, 0.238] | [0.419, 0.421] | [0.150, 0.150] | [0.099, 0.100] |  |

| **Model DtSt+** (LL: -10 597.1) |  |  |  |  |  |  |
| --- | --- | --- | --- | --- | --- | --- |
| | $\beta_1$ | $\beta_2$ | $\beta_3$ | $\rho$ | $\phi$ | $\theta$ |
| SI | [12.000, 14.000] | [0.200, 0.400] | [0.300, 0.500] | [0.120, 0.160] | [0.000, 0.050] | [0.300, 0.600] |
| TP | 12.657 | 0.238 | 0.420 | 0.150 | 0.100 | 0.500 |
| MLE | 12.658 | 0.235 | 0.429 | 0.156 | 0.135 | 0.473 |
| CI | [12.316, 12.998] | [0.219, 0.253] | [0.398, 0.463] | [0.15, 0.162] | [0.097, 0.173] | [0.441, 0.509] |

| **Model StSt+** (LL: $-11\,310.29 \pm 0.06$) | | | | | | |
| --- | --- | --- | --- | --- | --- | --- |
| | $\beta_1$ | $\beta_2$ | $\beta_3$ | $\rho$ | $\phi$ | $\theta$ |
| SI | [12.000, 14.000] | [0.200, 0.400] | [0.300, 0.500] | [0.120, 0.160] | [0.000, 0.050] | [0.300, 0.600] |
| TP | 12.657 | 0.238 | 0.420 | 0.150 | 0.100 | 0.500 |
| MLE | 12.794 | 0.235 | 0.407 | 0.149 | 0.102 | 0.467 |
| CI | [12.475, 13.099] | [0.222, 0.250] | [0.380, 0.440] | [0.144, 0.156] | [0.061, 0.143] | [0.437, 0.509] |

**Table 3:** Simulation study results for the models with starting interval (SI), true parameter (TP), maximum likelihood estimator (MLE) and calibration of the 95% confidence interval (CI).
| **Model St+St+** (LL: $-11\,102.11 \pm 0.13$) | | | | | | | |
| --- | --- | --- | --- | --- | --- | --- | --- |
| | $\beta_1$ | $\beta_2$ | $\beta_3$ | $\rho$ | $\phi$ | $\theta$ | $\sigma$ |
| SI | [12.000, 16.000] | [0.200, 0.400] | [0.300, 0.500] | [0.120, 0.160] | [0.000, 0.200] | [0.100, 0.500] | [0.001, 0.200] |
| TP | 12.657 | 0.238 | 0.420 | 0.150 | 0.100 | 0.300 | 0.050 |
| MLE | 12.390 | 0.248 | 0.431 | 0.152 | 0.113 | 0.313 | 0.034 |
| CI | [12.057, 12.684] | [0.235, 0.262] | [0.408, 0.460] | [0.145, 0.159] | [0.072, 0.163] | [0.291, 0.338] | [0.020, 0.052] |

### 3.3 Inference for rotavirus data

Parameter estimates as well as model diagnostics for the four models are given in Table 4. We report in the in the column ’coverage’ how often the actual data is covered by the pointwise 95% prediction interval of the respective model. For the stochastic transmission models StSt+ and St+St+ we also report how often the 95% prediction interval of the stochastic transmission model covers the data in order to investigate how much the observational noise additionally contributes to explaining the data. We find that matching Model DtSt to the data coincides surprisingly well with the analytical results for the susceptibility parameters, although, this model neither has an observational model nor a seasonal forcing component. However, the 95% prediction interval only covers 8.8% of the observed data and, hence, the model only poorly explains the variation in the data as is also reflected in a very low loglikelihood. Fitting the deterministic model with overdispersion (Model DtSt+) to the data improves the fit by several thousands log units. The 95% prediction interval of the estimated parameter values covers now 96.2% of the data.

**Table 4:**
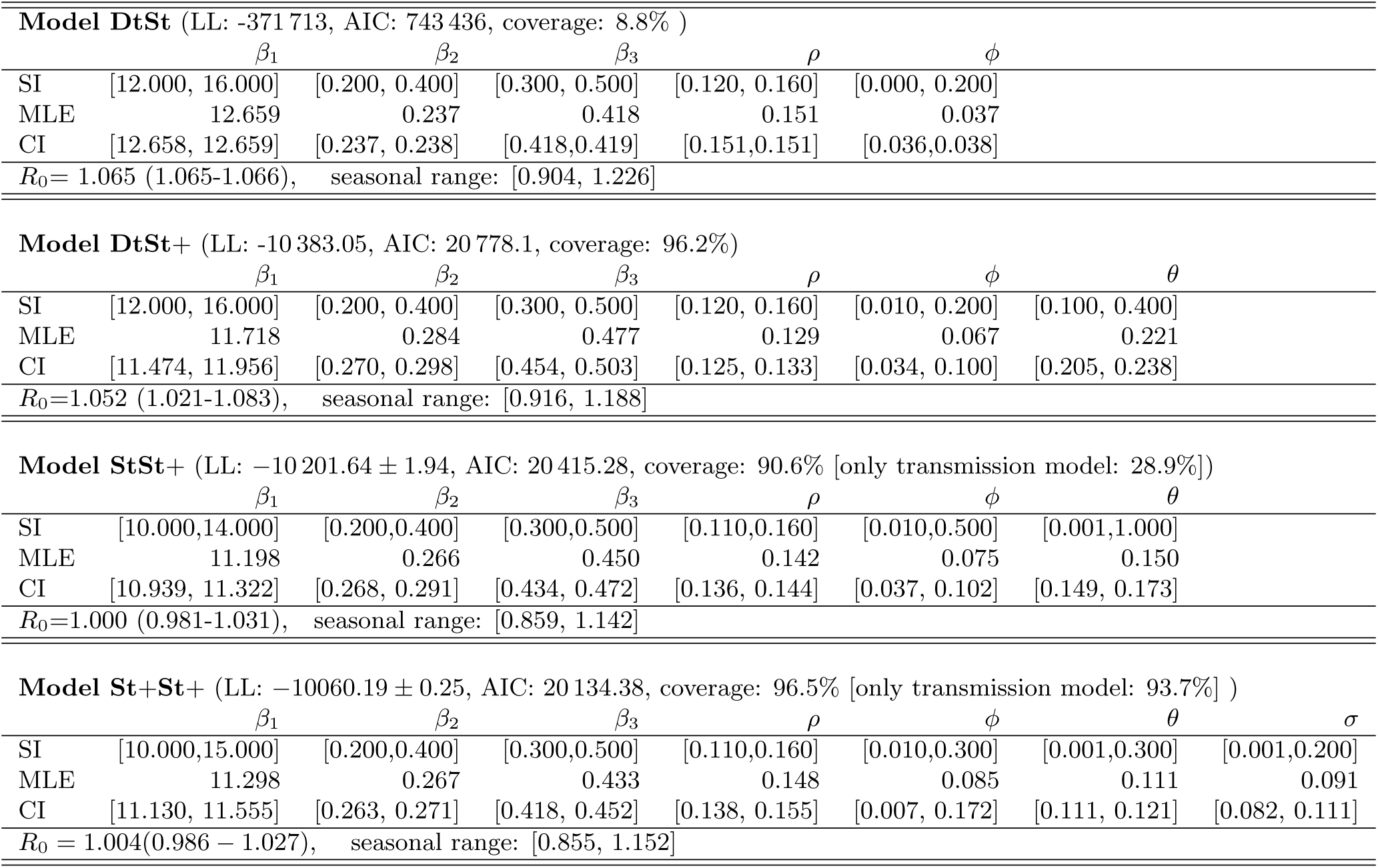
Inference results for the four models with starting interval (SI), maximum likelihood estimator (MLE), 95% confidence intervals (CI), basic reproduction number *R*_0_, 95% confidence interval for *R*_0_, seasonal variability of *R*_0_, coverage of the data by the 95 % prediction interval of the full model (coverage of the data 95 % prediction interval only generated by the transmission model), loglikelihood (LL), Akaike information criterion (AIC) and standard error of the Monte Carlo approximations.

| Model DtSt (LL: -371 713, AIC: 743 436, coverage: 8.8% ) |  |  |  |  |  |  |  |
| --- | --- | --- | --- | --- | --- | --- | --- |
| | $\beta_1$ | $\beta_2$ | $\beta_3$ | $\rho$ | $\phi$ | | |
| SI | [12.000, 16.000] | [0.200, 0.400] | [0.300, 0.500] | [0.120, 0.160] | [0.000, 0.200] |  |  |
| MLE | 12.659 | 0.237 | 0.418 | 0.151 | 0.037 |  |  |
| CI | [12.658, 12.659] | [0.237, 0.238] | [0.418,0.419] | [0.151,0.151] | [0.036,0.038] |  |  |
| $R_0= 1.065$ (1.065-1.066), seasonal range: [0.904, 1.226] | | | | | | | |
| Model DtSt+ (LL: -10 383.05, AIC: 20 778.1, coverage: 96.2%) |  |  |  |  |  |  |  |
| | $\beta_1$ | $\beta_2$ | $\beta_3$ | $\rho$ | $\phi$ | $\theta$ | |
| SI | [12.000, 16.000] | [0.200, 0.400] | [0.300, 0.500] | [0.120, 0.160] | [0.010, 0.200] | [0.100, 0.400] |  |
| MLE | 11.718 | 0.284 | 0.477 | 0.129 | 0.067 | 0.221 |  |
| CI | [11.474, 11.956] | [0.270, 0.298] | [0.454, 0.503] | [0.125, 0.133] | [0.034, 0.100] | [0.205, 0.238] |  |
| $R_0=1.052$ (1.021-1.083), seasonal range: [0.916, 1.188] | | | | | | | |
| Model StSt+ (LL: $-10\,201.64 \pm 1.94$, AIC: 20 415.28, coverage: 90.6% [only transmission model: 28.9%]) | | | | | | | |
| | $\beta_1$ | $\beta_2$ | $\beta_3$ | $\rho$ | $\phi$ | $\theta$ | |
| SI | [10.000,14.000] | [0.200,0.400] | [0.300,0.500] | [0.110,0.160] | [0.010,0.500] | [0.001,1.000] |  |
| MLE | 11.198 | 0.266 | 0.450 | 0.142 | 0.075 | 0.150 |  |
| CI | [10.939, 11.322] | [0.268, 0.291] | [0.434, 0.472] | [0.136, 0.144] | [0.037, 0.102] | [0.149, 0.173] |  |
| $R_0=1.000$ (0.981-1.031), seasonal range: [0.859, 1.142] | | | | | | | |
| Model St+St+ (LL: $-10060.19 \pm 0.25$, AIC: 20 134.38, coverage: 96.5% [only transmission model: 93.7%] ) | | | | | | | |
| | $\beta_1$ | $\beta_2$ | $\beta_3$ | $\rho$ | $\phi$ | $\theta$ | $\sigma$ |
| SI | [10.000,15.000] | [0.200,0.400] | [0.300,0.500] | [0.110,0.160] | [0.010,0.300] | [0.001,0.300] | [0.001,0.200] |
| MLE | 11.298 | 0.267 | 0.433 | 0.148 | 0.085 | 0.111 | 0.091 |
| CI | [11.130, 11.555] | [0.263, 0.271] | [0.418, 0.452] | [0.138, 0.155] | [0.007, 0.172] | [0.111, 0.121] | [0.082, 0.111] |
| $R_0= 1.004$(0.986 – 1.027), seasonal range: [0.855, 1.152] | | | | | | | |

The difference between Model DtSt+ and Model St+St+ is the nature of their transmission model and we find that a stochastic transmission model improves the fit by additional 180 log units. We find that the coverage of the 95% prediction interval decreases to 90.6%, however, the coverage of the prediction interval of the transmission model is 28.9%. The diagnostic plots (cf. Figures A.10 and A.12) show how the loglikelihood of the mif2 model and the parameters evolve with each iteration. In the diagnostic plot for Model StSt+ (Figure A.10) one can observe that after the loglikelihood of the perturbed model has increased significantly there is a small drop of the likelihood before stabilizing at a value which seems not optimal after having seen higher loglikelihood values at earlier iterations. However, re-running the particle filter at each iteration returned by the `mif2` model gives that the loglikelihood of the model of interest is increasing before stabilizing, cf. Figure 11. One possible explanation for the observed phenomenon is that the `mif2` model of Model StSt+ which includes extra variability in the parameters via random-walk perturbations explains the data better if the perturbations are larger. This indicates that the actual target model i.e. the one without perturbations, is not variable enough to explain the data well.

**Figure 4:**
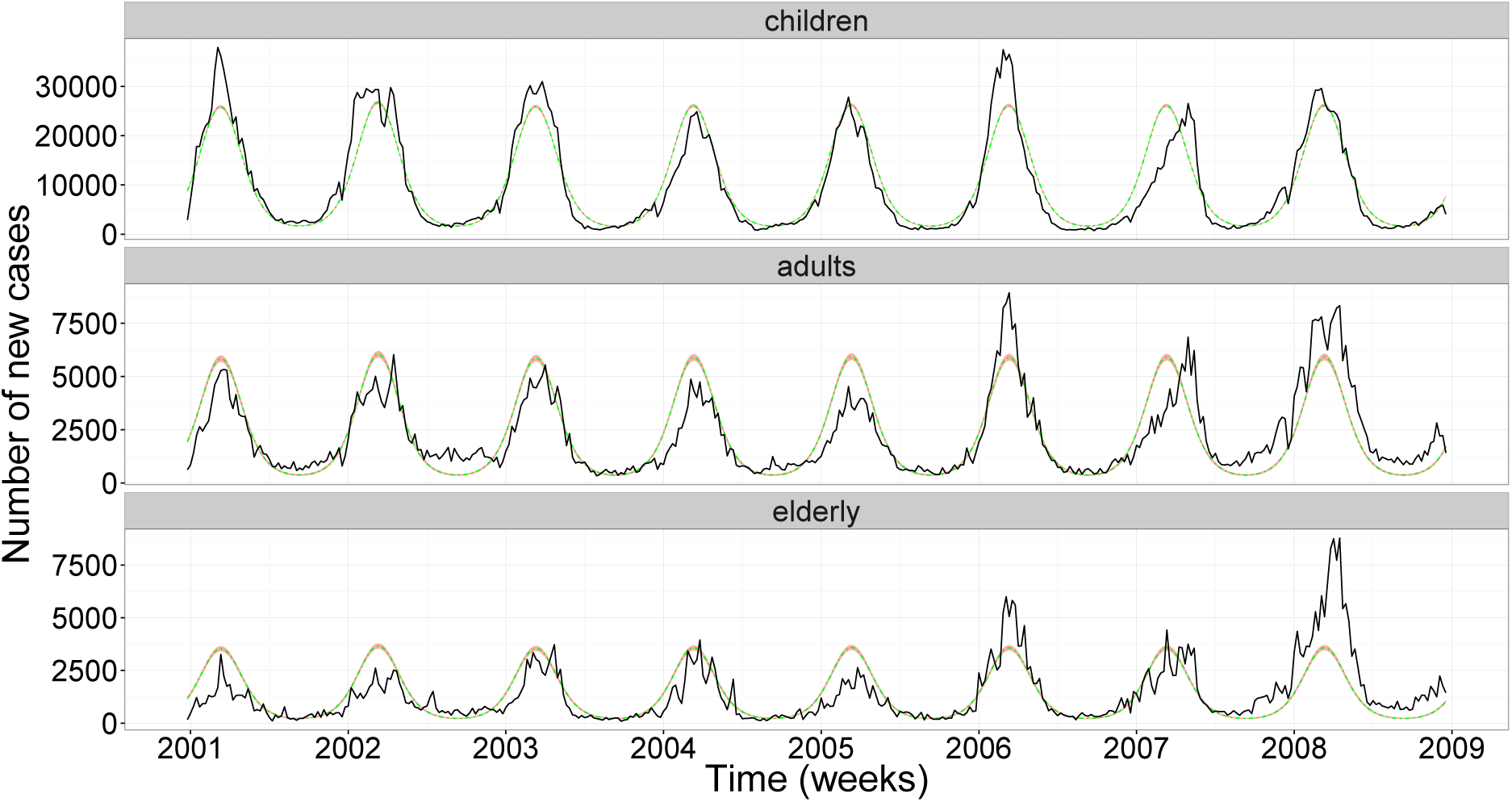
Model fit of Model DtSt to rotavirus incidence data (solid back line) and the model mean (solid light line). The 95% prediction interval is very thin so it is nearly indistinguishable from the model mean.

**Figure 5:**
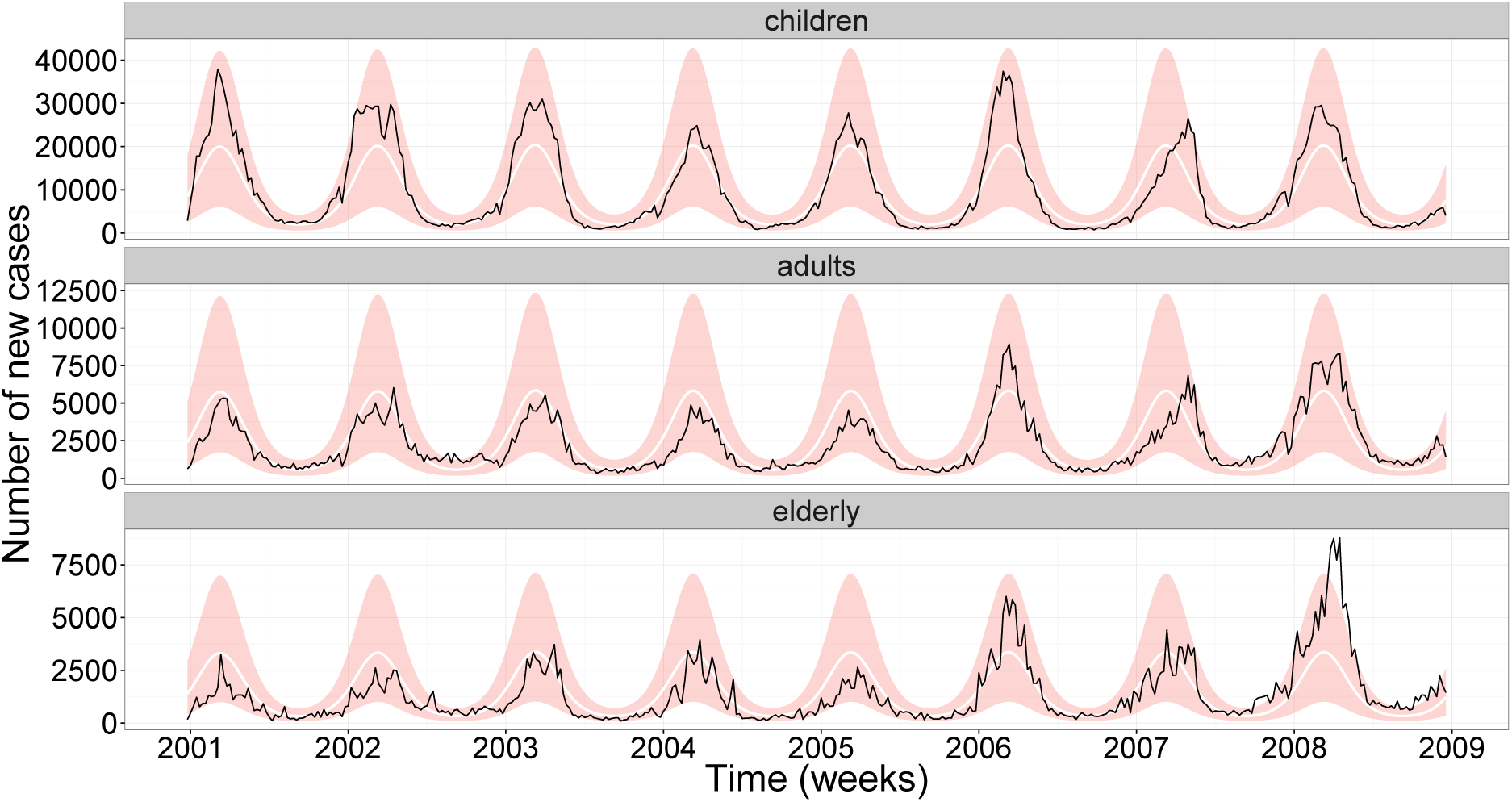
The 95% prediction interval (shading) for realizations of Model DtSt+ evaluated at the maximum likelihood estimator for the rotavirus incidence data (solid back line) and model mean (solid white line).

**Figure 6:**
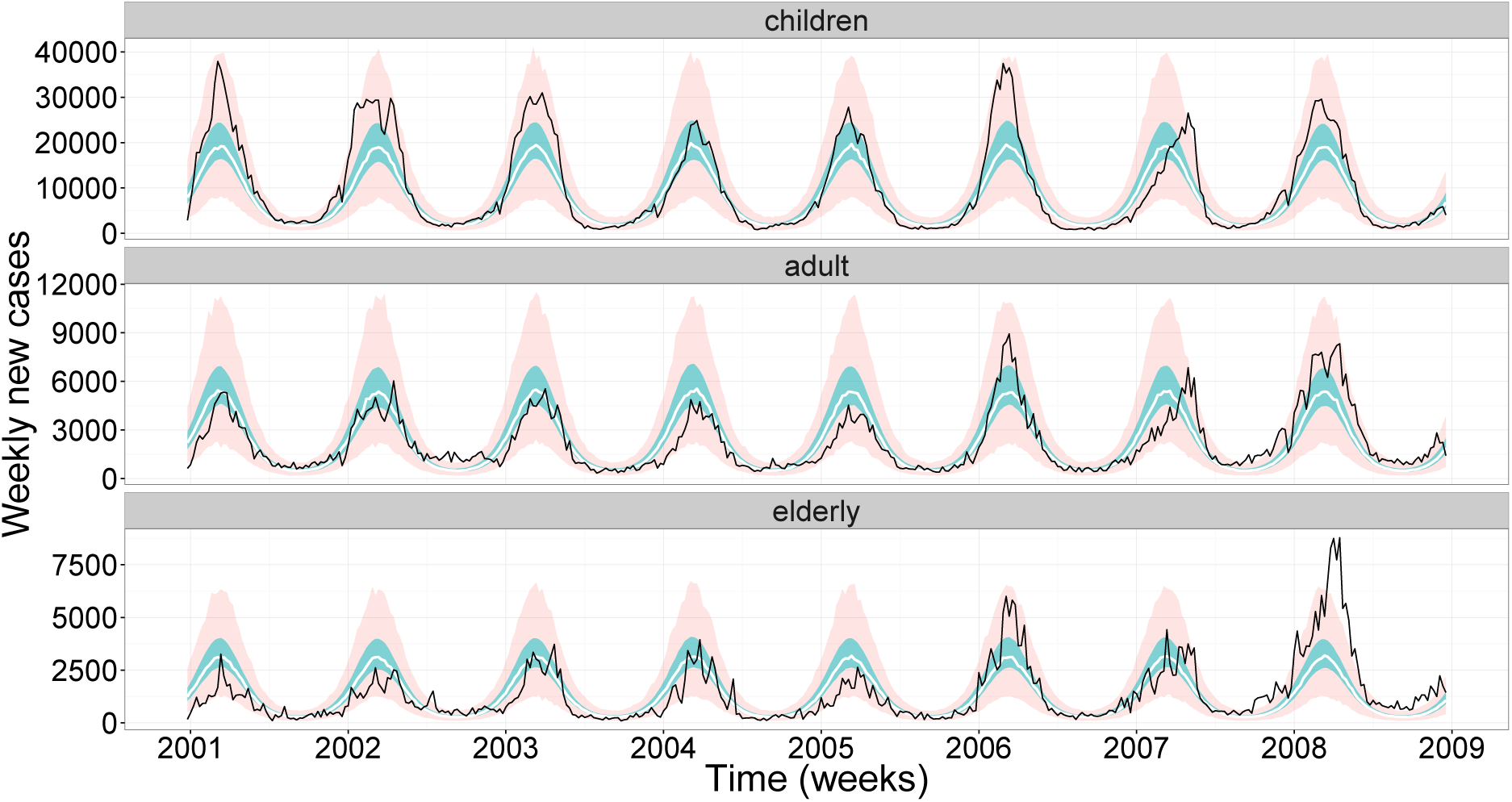
The 95% prediction interval (light shading) for 1000 realizations of Model StSt+ evaluated at the maximum likelihood estimator for the rotavirus incidence data (solid back line) and the median (solid white line). Furthermore, the 95 % prediction interval of these 1000 realizations for only the transmission model is shown (darker shading).

**Figure 7:**
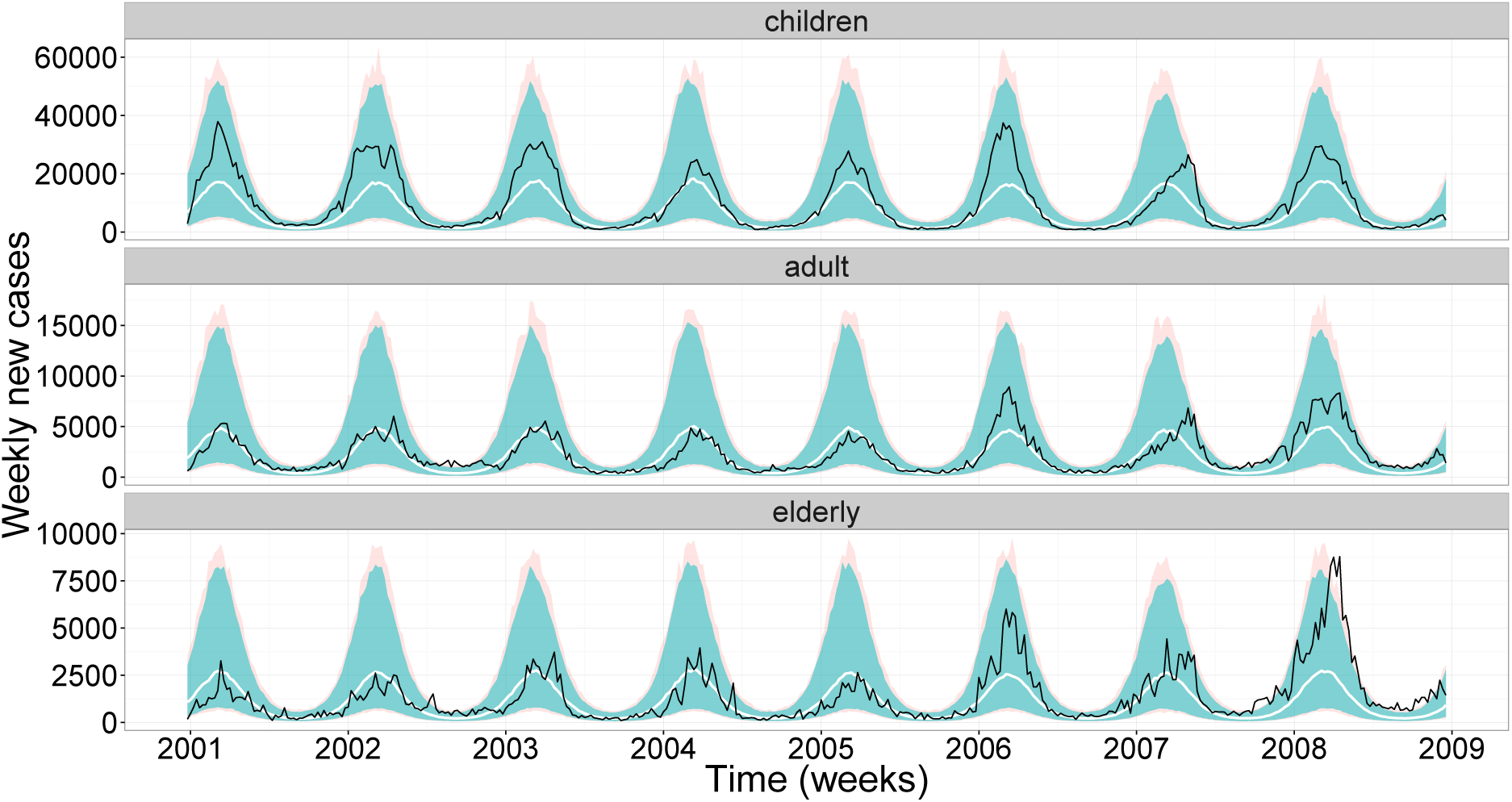
The 95% prediction interval (light shading) for 1000 realizations of Model St+St+ evaluated at the maximum likelihood estimator for the rotavirus incidence data (solid back line) and the median (solid white line). Furthermore, the 95 % prediction interval of these 1000 realizations for only the transmission model is shown (darker shading).

**Figure 8:**
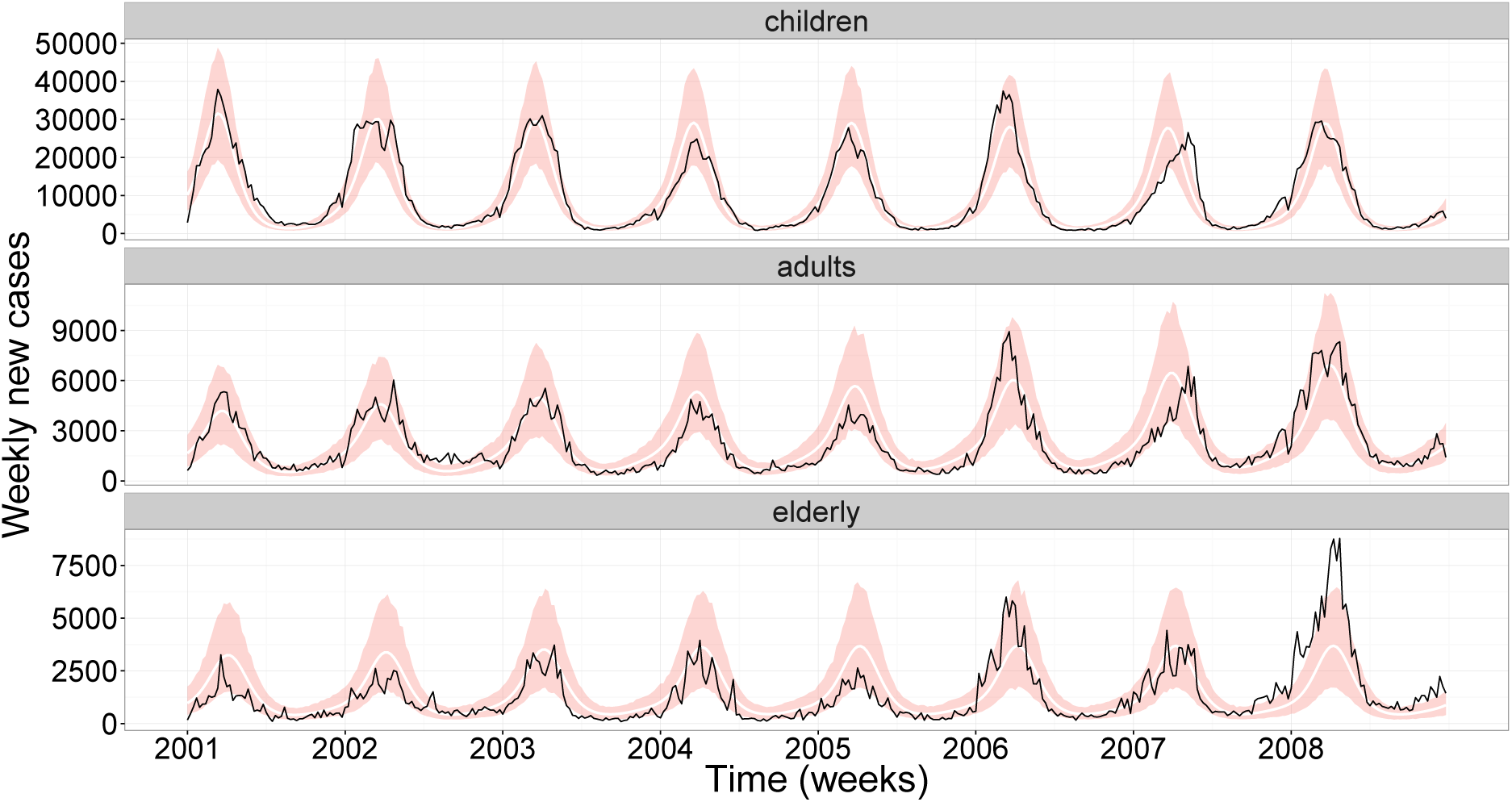
The 95% prediction interval (shading) for the model in Weidemann et al. (2013) assuming no underreporting, the model mean (solid white line) and the rotavrius incidence data (solid black line).

**Figure 9:**
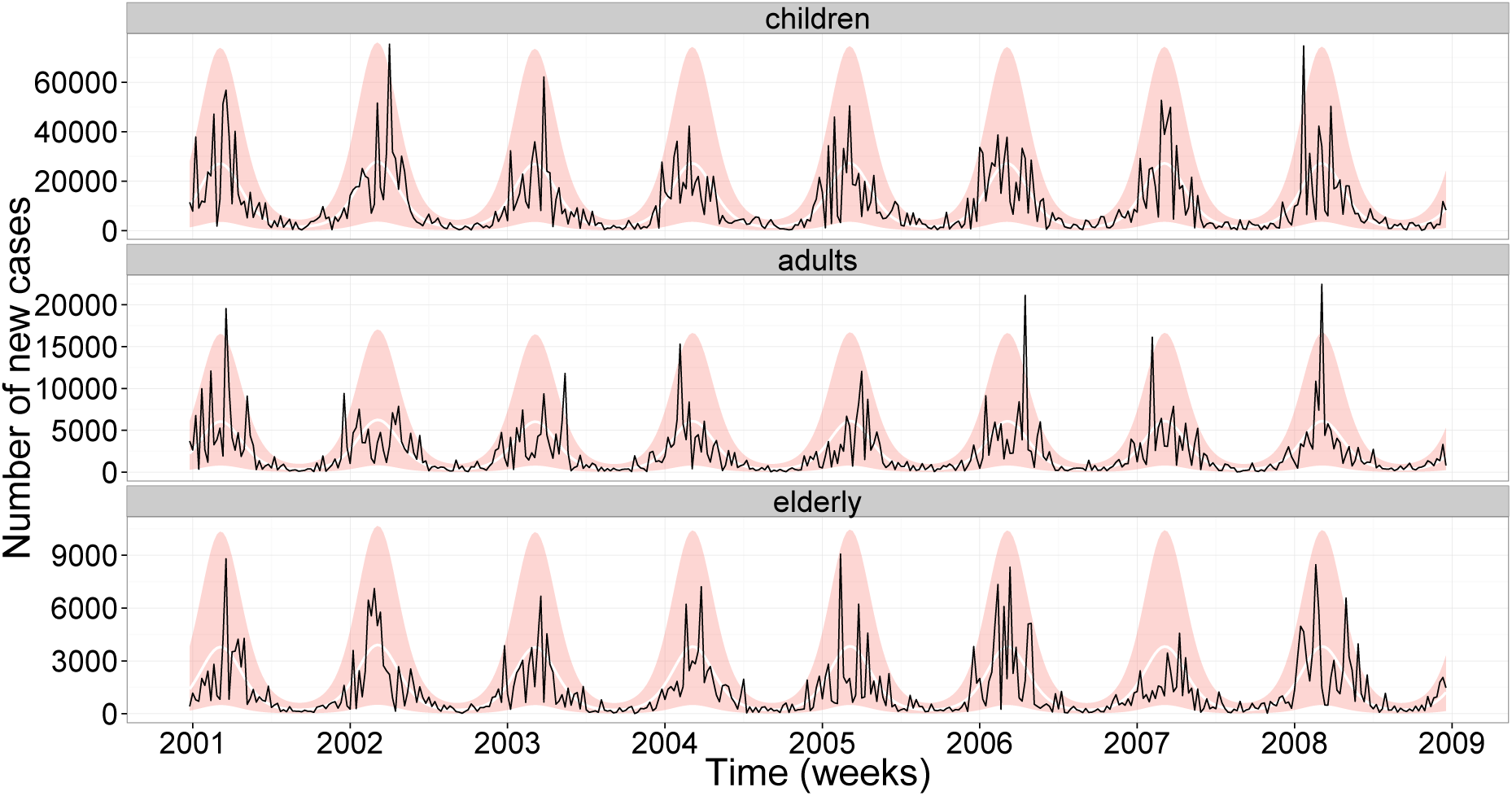
A Diagnostic plots. The 95% prediction interval (shading) for realizations of Model DtSt+ evaluated at the maximum likelihood estimator for simulated data (solid back line) and model mean (solid white line).

**Figure 10:**
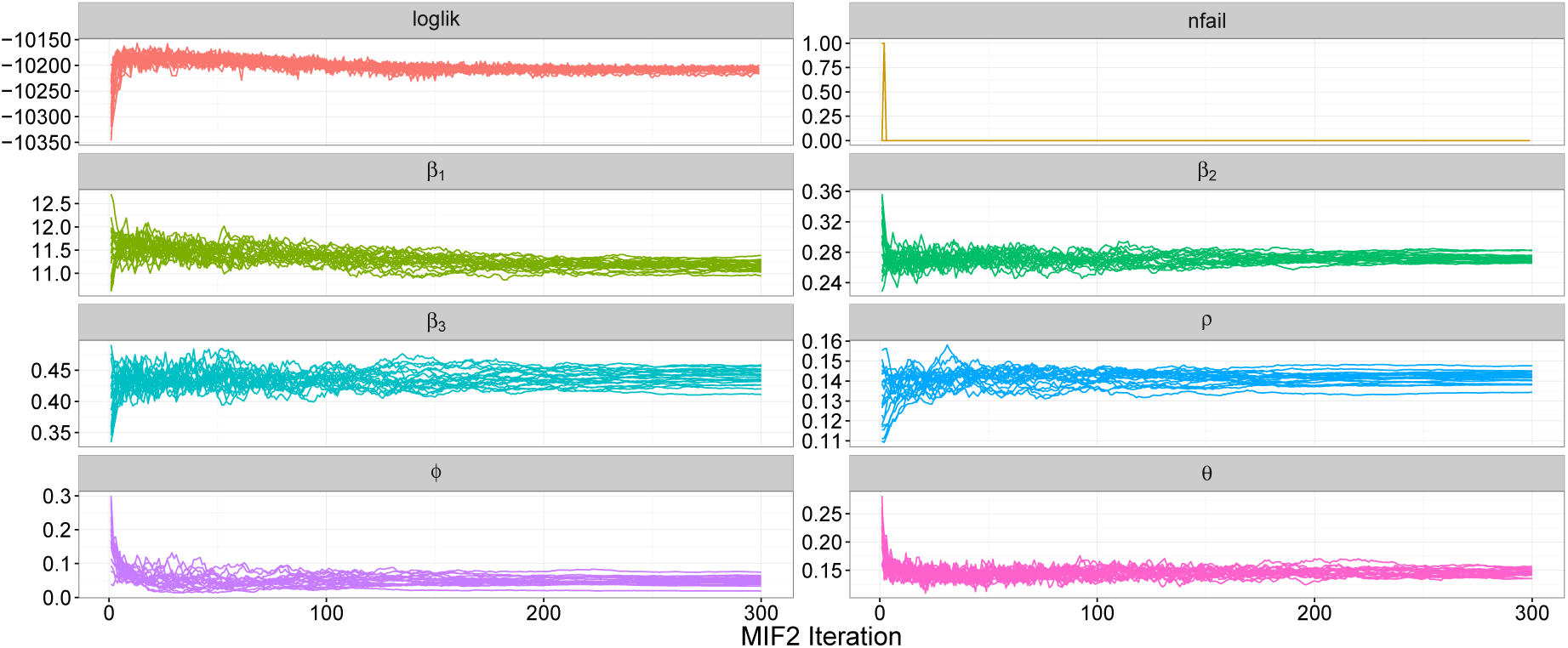
Diagnostic plot of the iterated filtering algorithm for Model StSt+for the rotavirus incidence data. Shown is the evolution of the loglikelihood and parameter estimates per `mif2` iteration for 20 trajectories with random starting values drawn from a hypercube.

**Figure 11:**
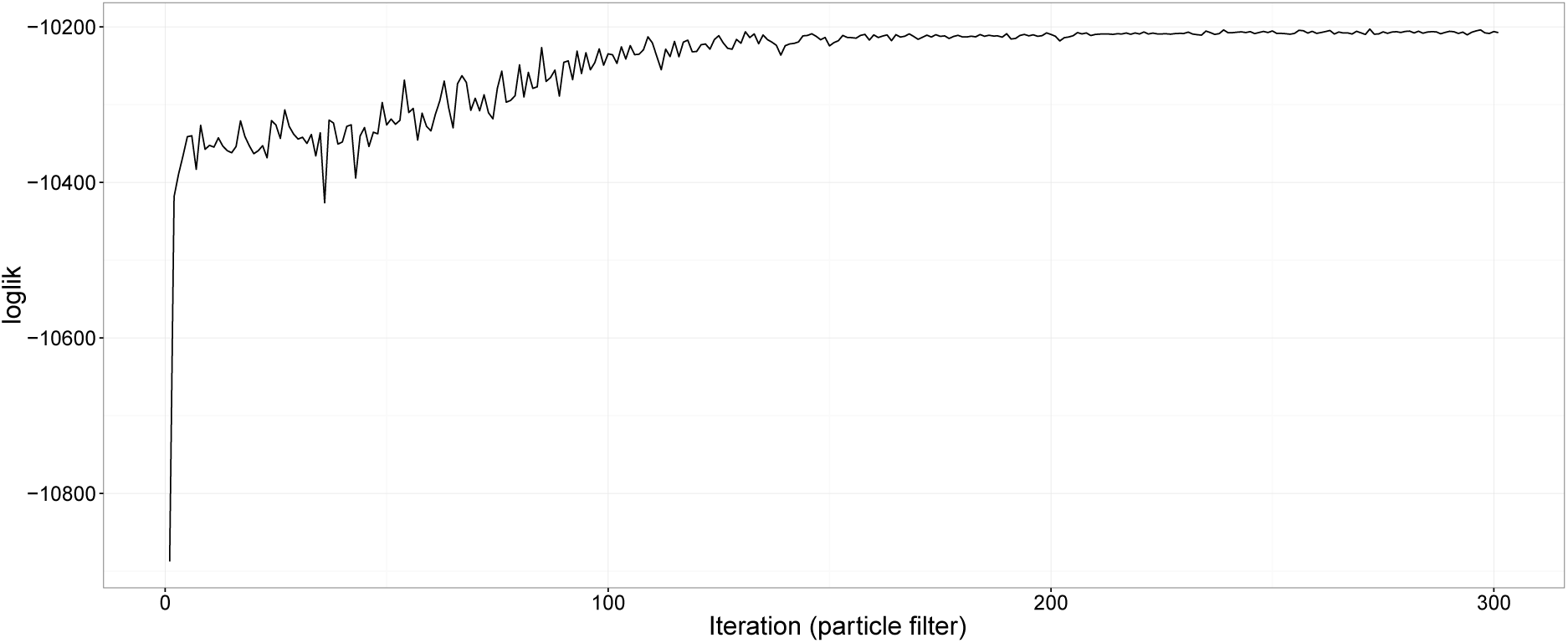
Particle filter evaluation for each iteration of one `mif2` run from Model StSt+ (Figure 10).

**Figure 12:**
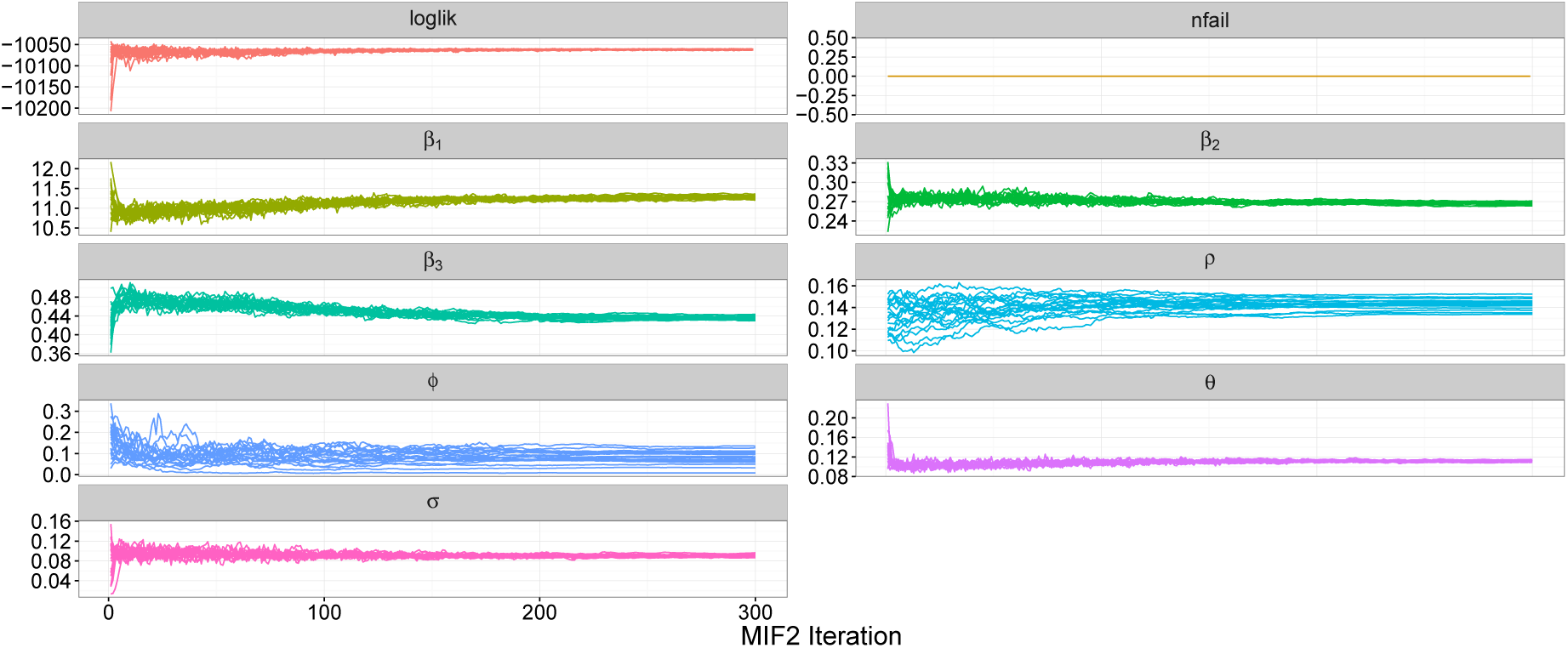
Diagnostic plot of the iterated filtering algorithm for Model St+St+ for the rotavirus incidence data. Shown is the evolution of the loglikelihood and parameter estimates per iteration for 20 trajectories with random starting values drawn from a hypercube.

This leads us to Model St+St+ which allows for additional variability by having stochastic transmission rates. It has the highest loglikelihood of the four models and is hence the best suited model to explain the data. The prediction interval covers 96.5% of the data. What is really interesting to note is that now the transmission model by itself is able to explain nearly all the data because its prediction interval covers 93.7%. This indicates that the observational noise component is not as strong as the three previous models suggested.

Although the data clearly lies in the 95% prediction interval for all models with overdispersion, we noticed that the model mean for the children in these models is slightly lower than the data for the first three years of our investigations. This does not occur for Poisson distributed observations so we initially presumed that the phenomenon might be an artifact of the negative binomial distribution. We carried out some additional analysis in order to understand this better. Firstly, we introduced age-dependent overdispersion parameters which improved the loglikelihood, however, did not raise the mean in the children compartment. Secondly, we assumed observations to be drawn from a left-censored normal distribution with the same mean and variance as for the negative binomial distribution in order to avoid the skewness of the negative binomial distribution. Also this approach did not raise the mean in the first age group so we conclude that the observed underestimation is not due to the negative binomial distribution but rather a general consequence of having overdipsersion in the model.

#### 3.3.1 Comparison with results from Weidemann et al. (2013)

We find that Model St+St+ explains the data very well, although the transmission component does not go into the same epidemiological detail as e.g. the work of Atkins et al. (2012) or Weidemann et al. (2013). Instead the model focuses on better capturing variability. Since we analysed the same data as in Weidemann et al. (2013) with the only difference that we directly scaled the data for under-reporting, it is very insightful to compare the model fits in order to see if it is worthwhile to model disease transmission very detailed and to give recommendations to future modellers of what we consider important. In Weidemann et al. (2013) a very detailed but deterministic transmission model with negative binomially distributed observations was used including 19 age classes, 3 susceptibility states, maternal antibody protection, distinguishing between symptomatic and asymptomatic cases resulting in an overall number of 266 age specific states. Moreover, the data was split into two data sets (former eastern (EFS) and western federal states (WFS)) so a region specific analysis was carried out and time specific birth and migration rates between age classes were included. In order to compare the fit of the models to the scaled up rotavirus data on equal grounds we slightly modified the inference sampling procedure which can be found on the Github repository indicated in Weidemann (2015). For the comparison we sampled 1000 times from the posterior distributions of the estimated parameters from their fit and solved the ODE system for the deterministic disease transmission model for each of the parameter constellations. From these solutions we then calculated the expected number of cases in each age group, respectively. Assuming that all region specific under-reporting rates were one (no decomposition of the expected number of incidences into the two regions) we then sampled the reported incidences according to a negative binomial observational distribution for each of the 1000 series and calculated the 2.5% and 97.5% sample quantiles. Note, that we have used the same dispersion parameter as in Weidemann et al. (2013) which is a fair approximation as long as the number of cases is fairly large. The plot of these prediction intervals and the rotavirus data used in our analysis is shown in Figure 8. The prediction bands for the resulting model are thinner than in Figure 7, however, if we as a measure of calibration calculate how many data points are covered by the prediction band we obtain 81.2%.

We conclude that a model which is simpler with respect to clinical detail in the disease transmission is sufficient to explain the rotavirus data as long as it accounts for structural and observational noise.

## 4 Discussion

In this paper we fitted four models, which differed with respect to the amount of stochasticity they allowed for, to routine rotavirus surveillance data from Germany. We demonstrated that a model which is simple with respect to rotavirus disease progression, but includes additonal population size independent variability in the underlying transmission model as well as over-dispersion in the observational model, is capable of explaining the available incidence data in a very satisfactory way. We also showed that a more detailed, but deterministic transmission model which only includes stochas-ticiticy as part of the observation process does not, from a fitting point of view, perform as well as the simpler model does. We find that structural noise in the transmission model clearly provides a better explanation of the data at hand. As stated in Bretó et al. (2009) such modelling is an appropriate way to “quantify the contributions of unknown and/or unmodelled processes” in the transmission model. We have evaluated our model fit with respect to how well the model is calibrated. Another method could be to assess the model’s sharpness by e.g. proper scoring rules (Gneiting and Raftery, 2007). Although our model was focusing on rotavirus transmission in Germany it could easily be modified to explain other infectious diseases which have comparable transmission characteristics such as influenza, chickenpox or pneumococcal disease. We conclude that an important modelling aspect for population based models is the adequate inclusion of variability: deterministic transmission models can be misleading because they hide model miss-specifications or unmodelled characteristics, which influence the disease transmission, in the observational noise component.

Throughout, we used a frequentist approach rather than a Bayesian setting for fitting the models. One advantage of this is that we did not have to assume priors for the unknown parameters. Such assumptions could obscure the fact that the models reach a complexity which makes individual parameters hardly identifiable and consequently the prior plays a crucial, but sometimes unintended, role. In this work we clearly address the question of identifiability and mathematically derive how many parameters can be estimated from the available data. Due to the simplicity of our model, every parameter is clearly understood and as a consequence we were able to estimate the basic reproduction number *R*_0_. However, it should be noted, that due to seasonal forcing this value is a yearly average, which changes the interpretation of *R*_0_ as a threshold value for an epidemic. Only if the yearly range does not cover the value one it can be interpreted in the traditional way. In our setting, the estimated *R*_0_ of all four models turned out to be close to one because the susceptibility parameter for children (*β*_1_) is pretty high while the ones for adults and elderly people (*β*_2_ and *β*_3_) are extremely low. This might have different reasons: first of all children might truly have a higher susceptibility to the disease than older individuals. Although our model indicated this, this might not be the most likely explanation. Secondly, it is possible that infants mix at a higher rate and expose themselves more to the disease by close body contact. Another possible scenario is that with higher age there might be partial immunity from earlier infections left which is why older individuals do not get ill. Furthermore, the severity of symptoms could play an important role for the reporting behaviour of the disease: it might be that symptoms are very severe (symptomatic cases) in infants which lead to a higher reporting rate, while cases in adults are less symptomatic or even asymptomatic and hence are not reported. However, even if symptoms are the same in all age classes, older individuals might not consult the doctor so often. Both of the last two explanations we would not detect, because we assumed the same scaling rate for the underreporting and fixed the rate of waning of immunity, *ω*, for all age classes. A sensitivity analysis for ω indicated that the longer the time of natural immunity last, the higher is *R*_0_. For example, if we choose the time of natural immunity 50 years, the basic reproduction rate increases to 1.44. Moreover, *R*_0_ depends heavily on the reporting rate we used to scale up the data- if the true underreporting rate is lower than we assumed, *R*_0_ increases. Of course we would have liked to disentangle all the previously mentioned explanations from each other, but the important message is that without any addition structural insights the available data is too coarse to do that. If the susceptibility parameters are not known the data cannot inform about the underreporting rate nor the duration of natural immunity which are necessary to calculate *R*_0_. We hence recommend to carry out studies specifically targeted to estimate these two quantities.

We have applied the `R package pomp` to rotavirus reporting data and gained helpful insights of how to handle this comprehensive package. It is a powerful package, however, inference is very time consuming and implementation requires non-intuitive model specific adjustments at times. More precisely, a major problem we faced initially was the high sensitivity of the estimation procedure with respect to starting values. We solved this problem by carrying out a detailed mathematically analysis investigating which parameters were actually identifiable. We were able to obtain an analytic solution for the susceptibility parameters of a simplified model, which we then used as a reasonable range for starting values. Furthermore, in order to make inference for the stochastic underlying transmission models feasible with respect to time, we had to accept an approximation error of the *τ*-leaping algorithm by choosing the simulation time interval rather big (1/10 week). The question arose if this approximation error is causing parts of the structural and observation noises. We carried out a sensitivity analysis for the simulation step size and decreased the step size to 1/40 weeks without finding a significant change of the noise parameters. Hence, we believe that the choice of the step size does not increase the structural and observational noise noticeably. Moreover, it turned out to be important to start the system well ahead of time so it could equilibrate before the actual observations start. Another valuable insight was that for a stochastic transmission model, Poisson distributed observations were not feasible because the model was simply not variable enough so the particle filter failed. Hence, overdispersion is a very important ingredient for successful estimation in the pomp model. It turned out that there is no precise rule of how to choose the magnitude of the random walk perturbations which the parameters undergo in the iterated filtering algorithm. It took some try and error to find reasonable values. Overall, the iterated filtering algorithm for the Models StSt+ and St+St+ took approximately 11h respectively when working with a computer with 20 cores. Therefore, we can highly recommend working with a computer cluster and parallelized `R` code as done by us. A comprehensive manual for seasonal incidence data can be found in Stocks (2017).

The reduced complexity of our model came at the cost of neglecting some potentially important clinical details of rotavirus transmission. We did not consider the short incubation time nor maternal immunity, we assumed that the waning immunity as well as the reporting rate, structural noise and dispersion parameter are age independent. Premature death was neglected and we did not use weekly demographic data or contact data such as the POLYMOD study (Mossong et al., 2008). Furthermore, we made the rather strong assumption that the time until an individual recovers as well as the time individuals stay in the different age classes is exponentially distributed. Despite this lack of detail the model still does its job and fits the data very well. Nevertheless, if the model is not only judged on the basis of model fit but is, e.g., used to investigate specific interventions such as vaccination strategies, more detailed models might be needed.

## Acknowledgements

We thank Aaron A. King for his very useful and detailed help with questions concerning the R package pomp. We furthermore thank Felix Weidemann for interesting discussions and sharing the code of his paper (Weidemann et al., 2013) and Gilles Kratzer for working on this topic in his master thesis. The computations were performed on resources provided by the Swedish National Infrastructure for Computing at PDC Centre for High Performance Computing. This manuscript was part of TS licentiate thesis so we thank the opponent Niel Hens and the examiner Martin Sköld for useful comments. TS and MH were in parts supported by the Swedish research council, grant number 2015_05182_VR.

## References

Althaus, C. L. (2014). Estimating the reproduction number of ebola virus (ebov) during the 2014 outbreak in West Africa. PLOS Currents: Outbreaks.

Anderson, R. M. and May, R. M. (1991). Infectious Diseases of Humans: Dynamics and Control. Oxford Science Publications.

Andersson, H. and Britton, T. (2000). Stochastic Epidemic Models and Their Statistical Analysis. Springer Lecture Notes in Statistics, 151.

Andrieu, C., Doucet, A., and Holenstein, R. (2010). Particle Markov chain Monte Carlo methods. Journal of the Royal Statistical Society, 72(3): 269–342.

Atchison, C., Lopman, B., and Edmunds, W. J. (2010). Modelling the seasonality of rotavirus disease and the impact of vaccination in England and Wales. Vaccine, 28(18):3118–3126.

Atkins, K. E., Shim, E., Pitzer, V. E., and Galvani, A. P. (2012). Impact of rotavirus vaccination on epidemiological dynamics in England and Wales. Vaccine, 30(3):552–564.

Bernstein, D. I. (March 2009). Rotavirus overview. The Pediatric Infectious Disease Journal, 3(3 Suppl):S50–3.

Bhadra, A., Ionides, E. L., Laneri, K., Pascual, M., Bouma, M., and Dhiman, R. (2011). Malaria in northwest India: Data analysis via partially observed stochastic differential equation models driven by Lévy noise. Journal of the American Statistical Association, 106(494):440–451.

Bretó, C., He, D., Ionides, E. L., and King, A. A. (2009). Time series analysis via mechanistic models. The Annals of Applied Statistics, 3(1):319–348.

CDC (Accessed December 4, 2016). http://www.cdc.gov/rotavirus/about/index.html.

Dennehy, P. H. (01.10.2015). Rotavirus infection: A disease of the past? Infectious Disease Clinics of North America, 29(4):617–635.

Diekmann, O., Heesterbeek, H., and Britton, T. (2013). Mathematical Tools for Understanding Infectious Disease Dynamics. Princeton Series in Theroretical and Computational Biology.

Doucet, A., de Freitas, N., and Gordon, N. (2001). Sequential Monte Carlo Methods in Practice. Springer-Verlag, New York.

Erhard, F., Friedel, C. C., and Zimmer, R. (2010). FERN-Stochastic Simulation and Evaluation of Reaction Networks. Systems Biology for Signaling Networks.

Freiesleben de Blasio, B., Kasymbekova, K., and Flem, E. (2010). Dynamic model of rotavirus transmission and the impact of rotavirus vaccination in Kyrgyzstan. Vaccine, 28(50):7923–3.

Fuchs, C. (2013). Inference for Diffusion Processes-With Applications in Life Sciences. Springer.

Gillespie, D. T. (1977). Exact stochastic simulation of coupled chemical reactions. The Journal of Physical Chemistry, 81(25):2340–2361.

Gillespie, D. T. (2001). Approximate accelatered stochastic simulation of chemically reacting systems. Journal of Chemical Physics, 115(4):1716–1733.

Gneiting, T. and Raftery, A. E. (2007). Strictly proper scoring rules, prediction, and estimation. Journal of the American Statistical Association, 102(477):359–378.

Grimmwood, K. and Lambert, S. B. (2009). Rotavirus vaccines: opportunities and challenges. Human Vaccines, 5(2):57–69.

He, D., Ionides, E. L., and King, A. A. (2009). Plug-and-play inference for disease dynamic: measles in large and small populations as a case study. Journal of the Royal Society Interface, 7(43):271–283.

Heesterbeek et al. (2015). Modeling infectious disease dynamics in the complex landscape of global health. Science, 347(6227):aaa4339.

Held, L. and Sabanés Bové, D. (2013). Applied Statistical Inference. Springer.

Ionides, E. L., Bhadra, A., Atchadé, Y., and King, A. A. (2011). Iterated filtering. The Annals of Statistics, 39(3):17761802.

Ionides, E. L., Bretó, C., and King, A. A. (2006). Inference for nonlinear dynamical systems. PNAS, 103(49):18438–18443.

Ionides, E. L., Nguyen, D., Atchadé, Y., Stoev, S., and King, A. (2015). Inference for dynamic and latent variable models via iterated, perturbed bayes maps. PNAS, 112(3):719–724.

Karlin, S. and Taylor, H. M. (1981). A Second Course in Stochastic Processes. Academic Press.

Keeling, M. J. and Rohani, P. (2008). Modeling Infectious Diseases in Humans and Animals. Princeton University Press.

Kendall, B. E., Briggs, C. J., Murdoch, W. W., Turchin, P., Ellner, S. P., McCauley, E., Nisbet, R. M., and Wood, S. N. (1999). Why do populations cycle? A synthesis of statistical and mechanistic modeling approaches. Ecology, 80(6):1789–1805.

King, A. A. and Ionides, E. L. (2016a). Iterated priciples: principles and practice. http://kingaa.github.io/short-course/mif/mif.html, [Accessed December 4, 2016].

King, A. A. and Ionides, E. L. (2016b). Likelihood for pomp models. http://kingaa.github.io/short-course/pfilter/pfilter.html, [Accessed December 4, 2016].

King, A. A., Ionides, E. L., Bret, C., Ellner, S., Kendall, B., Ferrari, M., Lavone, M., and Reuman, D. (2015). Introduction to pomp: Inference for partially-observed Markov processes. R vignette.

King, A. A., Nguyen, D., and Ionides, E. L. (2016). Statistical inference for partially observed Markov processes via the R package pomp. Journal of Statistical Software, 69(12):1–43.

Liu, J. and West, M. (2001). Combinig parameter and state estimation in simulation-based filtering. Sequential Monte Carlo Methods in Practice, pages 197–223.

Martinez, P. P., King, A. A., Yunusd, M., Faruqued, A. S. G., and Pascual, M. (2016). Differential and enhanced response to climate forcing in diarrheal disease due to rotavirus across a megacity of the developing world. PNAS, 113(15):4092–4097.

Martinez-Bakker, M., King, A. A., and Rohani, P. (2015). Unraveling the transmission ecology of polio. PLOS Biology, 13(6):e1002172.

May, R. M. (2004). Uses and abuses of mathematics in biology. Science, 303(5659):790–793.

Mossong, J., Hens, N., Jit, M., Beutels, P., Auranen, K., Mikolajczyk, R., Massari, M., Salmaso, S., Scalia Tomba, G., Wallinga, J., Heijne, J., Sadkowska-Todys, M., Rosinska, M., and Edmunds, W. J. (2008). Social contacts and mixing patterns relevant to the spread of infectious diseases. PLOS Medicine, 5(3):e74.

Nelder, J. A. and Mead, R. (1965). A simplex method for function minimization. The Computer Journal, 7(4):308–313.

O’Neill, P. D. (2010). Introduction and snapshot review: relating infectious disease transmission models to data. Statistics in Medicine, 29(20):2069–2077.

Pitzer, V. E., Viboud, C., Simonsen, L., Steiner, C., Panozzo, C. A., Alonso, W. J., Miller, M. A., Glass, R. I., Glasser, J. W., Parashar, U. D., and Grenfel, B. T. (2009). Demographic variability, vaccination, and the spatiotemporal dynamics of rotavirus epidemics. Science, 325(5938):290–294.

Pitzer et al. (2012). Direct and indirect effects of rotavirus vaccination: Comparing predictions from transmission dynamic models. PLOS one, 7(8):e42320.

Press, W. H., Flannery, B. P., Teukolsky, S. A., and Vetterling, W. T. (2007). Numerical Recipes: The Art of Scientific Computing (3rd edition). Cambridge University Press.

RKI (Accessed December 4, 2016). Survstat. https://survstat.rki.de/.

Robert, C. P. and Casella, G. (2004). Monte Carlo Statistical Methods. 2nd editions. Springer-Verlag.

Shim, E., Banks, C., and Castillo-Chavez, H. T. (2006). Seasonality of rotavirus infection with its vaccination. AMS Contemporary Mathematics Book Series, 410.

Statistisches Bundesamt (Accessed December 4, 2016). https://www.destatis.de/DE/ZahlenFakten/GesellschaftStaat/Bevoelkerung/Bevoelkerungsstand/Tabellen_/lrbev01.html.

Stocks, T. (2017). Github account. https://github.com/theresasophia, [Accessed March 10, 2017].

Sugihara, G. and May, R. M. (1990). Nonlinear forecasting as a way of distinguishing chaos from measurement error in time series. Nature, 344(6268):734–741.

The World Bank (Oct, 2016). http://data.worldbank.org/indicator/sp.dyn.le00.in?locations=de.

Toni, T., Welch, D., Strelkowa, N., Ipsen, A., and Stumpf, M. P. (2009). Approximate Bayesian computation scheme for parameter inference and model selection in dynamical systems. Journal of the Royal Society, 6(31).

Van Effelterre, T., Soriano-Gabarr, M., Debrus, S., Claire Newbern, E., and Gray, J. (2010). A mathematical model of the indirect effects of rotavirus vaccination. Epidemiology and Infection, 138(6):884–897.

Weidemann, F. (2015). Bayesian Inference for Infectious Disease Transmission Models Based on Ordinary Differential Equations. https://edoc.ub.uni-muenchen.de/19060/1/weidemann_felix.pdf, [Accessed December 4, 2016], Ludwig-Maximilians-University Munich.

Weidemann, F., Dehnert, M., Koch, J., Wichmann, O., and Höhle, M. (2013). Bayesian parameter inference for dynamic infectious disease modeling: rotavirus in Germany. Statistcs in Medicine, 33(9):1580–1599.

Weidemann, F., Dehnert, M., Koch, J., Wichmann, O., and Höhle, M. (2014). Modelling the epidemiological impact of rotavirus vaccination in Germany - a Bayesian approach. Vaccine, 32(40):5250–5257.

Wood, S. N. (2010). Statistical inference for noisy nonlinear ecological dynamic systems. Nature, 466: 1102–1104.

